# Nuclear lamin A/C phosphorylation by loss of Androgen Receptor is a global determinant of cancer-associated fibroblast activation

**DOI:** 10.1101/2023.06.28.546870

**Authors:** Soumitra Ghosh, Jovan Isma, Luigi Mazzeo, Annagiada Toniolo, Christian Simon, G. Paolo Dotto

## Abstract

Alterations of nuclear structure and function, and associated impact on gene transcription, are a hallmark of cancer cells. Little is known of these alterations in Cancer-Associated Fibroblasts (CAFs), a key component of the tumor stroma. Here we show that loss of androgen receptor (AR), which triggers early steps of CAF activation in human dermal fibroblasts (HDFs), leads to nuclear membrane alterations and increased micronuclei formation, which are unlinked from induction of cellular senescence. Similar alterations occur in fully established CAFs, which are overcome by restored AR function. AR associates with nuclear lamin A/C and loss of AR results in a substantially increased lamin A/C nucleoplasmic redistribution. Mechanistically, AR functions as a bridge between lamin A/C with the protein phosphatase PPP1. In parallel with a decreased lamin-PPP1 association, AR loss results in a marked increase of lamin A/C phosphorylation at Ser 301, which is also a feature of CAFs. Phosphorylated lamin A/C at Ser 301 binds to the transcription promoter regulatory region of several CAF effector genes, which are upregulated due to the loss of AR. More directly, expression of a lamin A/C Ser301 phosphomimetic mutant alone is sufficient to convert normal fibroblasts into tumor-promoting CAFs of the myofibroblast subtype, without an impact on senescence. These findings highlight the pivotal role of the AR-lamin A/C-PPP1 axis and lamin A/C phosphorylation at Ser 301 in driving CAF activation.

## INTRODUCTION

Alterations in nuclear structure and function are characteristic of tumor cells and are often used as prognostic and diagnostic markers ^1–3^. However, the existence and functional significance of nuclear alterations in surrounding stromal cells and specifically cancer associated fibroblasts (CAFs) have yet to be explored. CAFs can play a key role at all stages of cancer development. They consist of heterogeneous populations of cells with distinct tumor enhancing properties ^4–11^ and, at least in skin, display increased genomic instability and specific genomic alterations conferring selective advantages for cancer-stromal cells co expansion ^12, 13^.

Mammalian cells express two types of nuclear lamins, which are key components of the scaffold around the nucleus ^14, 15^. Nuclear lamins play an essential role at the level of the nuclear envelope and inside the nucleus, impacting on transcription, splicing, chromatin organization and DNA repair ^16, 17^. While type B lamins can play an important role in many aspects of cell physiology, most studies have focused on intranuclear localization and function of type A/C lamins ^14, 16, 18^. While nuclear lamins have been highly studied in the context of aging and cancer ^16^, to our knowledge, their role in CAFs has not been investigated.

A close interconnection exists between the senescence of stromal fibroblasts and induction of a Senescence Associated Secretory Phenotype (SASP), consisting of the production of pro-tumorigenic cytokines, matrix components and remodeling enzymes ^19, 20^. We have previously shown that the androgen receptor (AR) plays a key role in the control of early steps of CAF activation, being involved in the concomitant transcriptional repression of senescence (*CDKN1A*) and SASP effector genes ^21^. AR expression is down-modulated in dermal fibroblasts of the intact skin *in vivo* as the result of photoaging and direct UVA exposure and is associated with early stages of skin cancer development, specifically actinic keratosis lesions, which develop in >80% of the aging Caucasian populations ^21^.

Fully established CAFs exhibit the expression of SASP genes but have evaded molecular mechanisms of cellular senescence and are capable of co-evolving and growing alongside cancer cells ^22^. The identification of mechanism(s) that are specifically involved in CAF activation of various subtypes without having an impact on senescence would be of substantial interest to prevent the entire cancer process.

Here we show that AR loss, by either genetic or pharmacological means, causes significant nuclear alterations in HDFs unlinked from p53-dependent cellular senescence, with similar abnormalities occurring in CAFs, which can be overcome by restored AR function. AR physically associates with lamin A/C and is a determinant of its proper localization at the nuclear membrane versus chromatin compartment. AR loss compromises the association of lamin A/C with the nuclear protein phosphatase PPP1 and results in increased lamin A/C phosphorylation at Ser 301, which is also a distinguishing feature of CAFs. Phospho-301 lamin A/C is recruited to the promoter / transcription regulatory region of CAF effector genes of the myofibroblast type and a phospho-mimetic (Ser/Asp) 301 lamin A/C is sufficient to induce expression of these genes and trigger CAF activation without an impact on senescence. Thus, nuclear abnormalities and lamin A/C phosphorylation at a specific site are CAF features of functional significance resulting from the loss of AR.

## RESULTS

### 1. Nuclear membrane integrity depends on sustained AR expression in normal dermal fibroblasts

Loss of androgen receptor (AR) in human dermal fibroblasts (HDFs) triggers early steps of CAF activation, with induction of p53-dependent cellular senescence and expression of multiple CAF effector genes ^21^. Lentiviral-mediated silencing of the *AR* gene in three different strains of HDFs resulted in consistent nuclear alterations, with irregular shape and decreased circularity index, membrane blebbing and micronuclei (MN) formation (collectively referred to as “nuclear abnormalities„) (Fig.1A-C; Fig. S1A,B). Transmission electron microscopy (TEM) of *AR* silenced cells showed nuclear envelope invaginations, nuclear blebbing, and micronuclei formation, which were not detectable in control cells (Fig. 1D, Sup. Fig. S1C).

**Figure 1.**
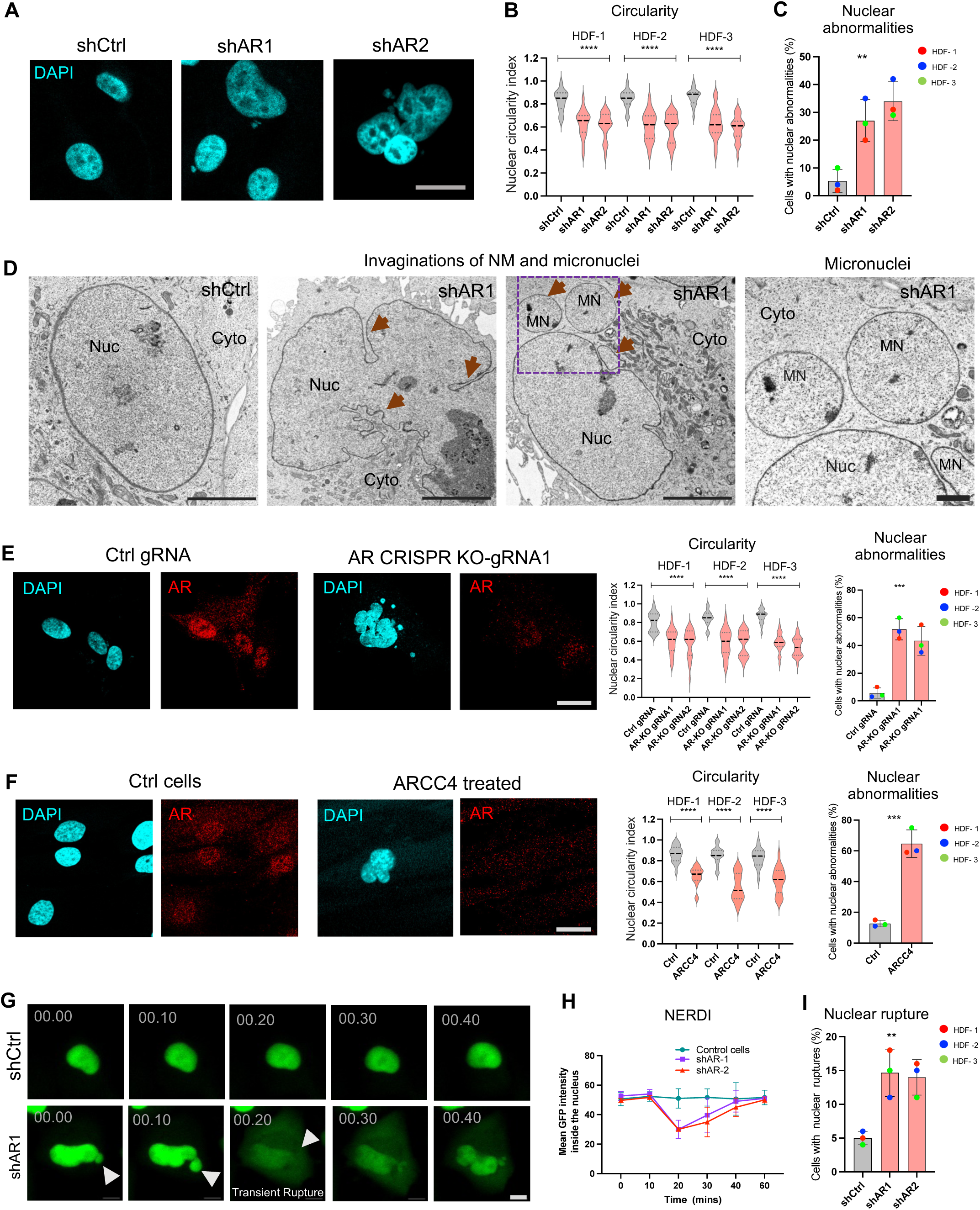
Loss of AR in HDFs leads to nuclear membrane abnormalities and rupture. A-C) Alterations of nuclear morphology in three primary HDF strains infected with two *AR* silencing lentiviruses (shAR1 and shAR2) versus control (shCtrl). A): representative images of DAPI-stained nuclei at 6 days after infection. Scale bar: 10 μm. Knockdown of *AR* expression is shown in Fig. S1A,B. B) Quantifications of nuclear circularity index, measured on central sections of DAPI-stained nuclei by the ImageJ particle analysis tool. Shown are violin plots of individual nuclei values for each of the three HDF strains plus/minus *AR* silencing (>100 cells per condition). Central dotted line indicates median value and two additional dotted lines indicate the 25th and 75th percentiles. n= 100 cells, ****P<0.0001, non-parametric one-way ANOVA Kruskal–Wallis test. C) Percentages of cells with nuclear abnormalities (micronuclei and nuclear blebbing) determined by inspection of digitally acquired images of the three HDF strains plus/minus *AR* silencing (>100 cells per condition). n=3 strains, **P<0.01; non-parametric one-way ANOVA Kruskal–Wallis test, mean±SE. D) Transmission electron microscopy images of nuclei from control and *AR*-silenced HDFs. Shown are representative images, with arrowheads indicating nuclear membrane invaginations and micronuclei. Scale bars: 5 μm (three left panels) and 1 μm (right panel). Additional images are shown in Fig. S1C. E) Alteration of nuclear morphology in three HDF strains with CRISPR-mediated deletion of *AR*, visualized by fluorescence confocal microscopy. HDFs were infected with high titer lentiviruses expressing either scrambled gRNA (Ctrl gRNA) or two independent *AR*-targeting gRNAs (gRNA1 and gRNA2). *AR* gene targeting and loss of AR expression are shown in Fig. S1F,G. Shown are representative images of DAPI stained nuclei and AR immunostaining of HDFs infected with control versus *AR*-targeting CRISPR-lentiviruses together with quantification of nuclear circularity and nuclear abnormalities as in (B) and (C). For circularity, n=>100 cells per condition; for abnormalities, n=3 HDF strains;****P<0.0001; ***P<0.001; non-parametric one-way ANOVA Kruskal–Wallis test, mean±SE. Scale bar: 10 μm. F) Nuclear alterations in three HDF strains plus/minus treatment with the AR degrading ARCC4 compound. Cells were treated with either EtOH solvent control (Ctrl) or ARCC4 (1 μM) for 72h followed by DAPI and anti-AR antibody staining. Shown are representative images together with quantification of nuclear circularity and nuclear abnormalities as in the previous panels. For circularity, n=>50 cells per condition; for abnormalities, n=3 strains; ****P<0.0001, ***P<0.001; paired T-test, mean±SE. Scale bar: 10 μm. G) Representative snapshots of time-lapse imaging analysis of HDFs plus/minus *AR* gene silencing and expression of a green fluorescence protein with nuclear localization signal (GFP–NLS) for dynamic assessment of nuclear envelope integrity. GFP-NLS expressing HDFs were infected with a control versus *AR* silencing lentiviruses followed, six days later, by seeding on glass-bottom chamber slides, for time-lapse imaging. Images were acquired for 24 hours at 10 minutes intervals. Images show a transient nuclear envelope rupture during interphase (NERDI) event in HDFs with silenced *AR*, with arrowheads pointing to a nuclear bleb formation (left two images) and subsequent transient nuclear rupture (middle image). Gradual recovery of GFP-NLS into the nucleus is shown in last two images. Time is shown on the top of each panel (hours: minutes). Scale bar: 5 μm. H) Kinetics of NERDI as extracted from GFP–NLS live-cell imaging of control and shAR HDF cells, with 10-min intervals. The graph indicates data obtained from the mean nuclear GFP signal intensity of 20 cells from each condition in 3 independent strains. I) Quantifications of the percentage of cells with nuclear envelope rupture in shCtrl, shAR1, shAR2 conditions. **P<0.01; non-parametric one-way ANOVA Kruskal–Wallis test, n=3 HDF strains. The data shown are the mean ± SE.

Similar nuclear abnormalities were observed in *AR* silenced HDFs cultured on collagen-coated soft surfaces (Fig. S1D) and upon *AR* silencing in p53-knockdown HDFs (Fig. S1E), indicating that the consequences of AR loss on nuclear structural integrity do not depend on substrate stiffness condition nor are linked to p53-dependent cellular senescence.

We sought validation of the results by deletion of the *AR* gene by CRISPR/Cas9 technology ^23^. Given the negative effects of *AR* loss in HDFs ^21^, we resorted to infection of three different strains with high titer lentiviruses co-expressing Cas9 together with two different *AR*-targeting guide RNAs (gRNAs), followed by analysis of pooled polyclonal cell populations shortly after infection. Surveyor assays ^24^ and immunofluorescence staining with anti-AR antibodies indicated that the *AR* gene was targeted and AR protein expression was effectively reduced by this approach in the majority of the cells (Fig. S1F,G). Combined immunofluorescence and nuclear morphometric analysis showed that CRISPR-targeting of *AR* in HDFs resulted in similar nuclear alterations as upon shRNA-mediated gene silencing (Fig. 1E).

As an alternative pharmacological approach, HDFs were treated with two different AR inhibitors, one, ARCC4, causing PROTAC-mediated degradation of AR ^25^ and the other, AZD3514, suppressing AR activity through both ligand competitive and non-competitive mechanisms ^26^. Immunofluorescence analysis showed that treatment of multiple HDF strains with either inhibitor caused effective loss of AR expression that was accompanied by drastically increased nuclear alterations (Fig. 1F, Fig. S1H).

The maintenance of nuclear membrane integrity is a dynamic process ^27^. Time-lapse fluorescence microscopy of HDFs expressing a green fluorescent protein fused with a nuclear localization signal (GFP-NLS) revealed a highly dynamic pattern of nuclear blebbing in HDFs with the silenced *AR* gene, associated with transient nuclear envelope rupture during interphase (NERDI) ^27^. A rapid decrease of nuclear GFP signal was observed in these cells that was followed by a gradual accumulation back into the nucleus upon nuclear membrane repair ^27^ (Fig. 1G,H, Fig. S1I). Permanent nuclear rupture was also observed in some of the cells (Fig.1I, Fig.S1I).

Thus, loss of AR expression in normal dermal fibroblasts results in compromised nuclear membrane integrity.

### 2. Nuclear abnormalities are an AR-dependent feature of CAFs

Decreased *AR* expression is a hallmark of CAFs, with CAF activation being counteracted by increased AR expression ^21^. Nuclear changes comparable to those resulting from *AR* loss in HDFs were found in a panel of skin Squamous cell carcinoma (SCC)-derived CAFs compared to matched HDFs from flanking unaffected skin of the same patients (Fig. 2A, Fig. S2A). Overexpression of *AR* in the CAFs rescued their nuclear abnormalities, linking altered nuclear structure in these cells to levels of *AR* expression (Fig. 2B, Fig. S2B). AR can positively regulate its own expression as part of a positive feedback regulatory loop ^28^. Treatment of CAFs with the AR agonist ostarine ^29^ resulted in a concomitant increase of endogenous AR expression and suppressed nuclear abnormalities (Fig. 2C,D; Fig. S2C)

**Figure 2.**
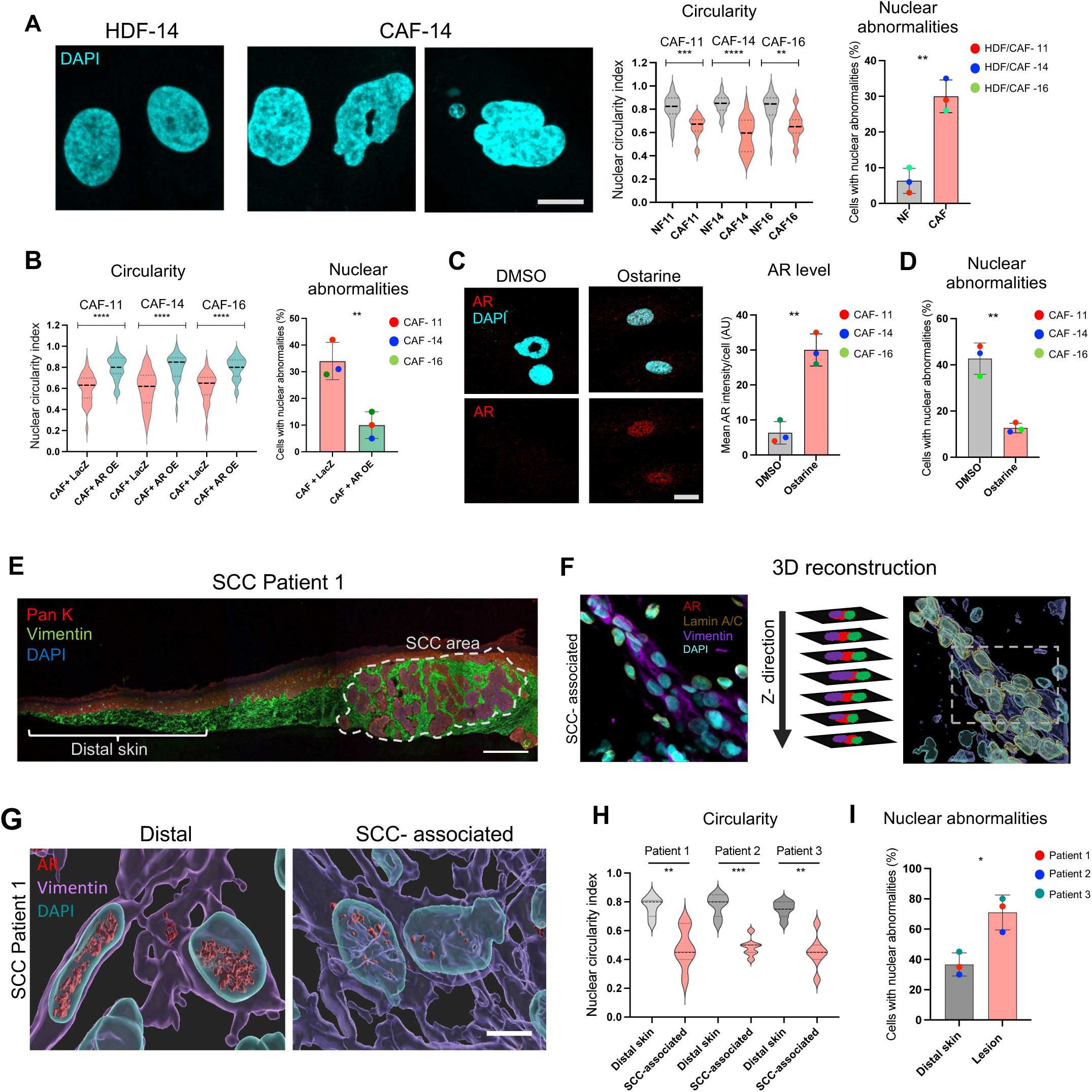
Nuclear abnormalities in cancer-associated fibroblasts (CAFs). (A) Left, representative confocal images of DAPI-stained nuclei of CAFs and matched HDFs isolated from the same patients. Scale bar: 10 µm. Right, quantification of nuclear circularity index and nuclear abnormalities in three pairs of CAFs and matched HDFs as in Fig. 1B,C. For circularity, n=>100 cells per condition; for abnormalities, n=3 HDF strains; ****P<0.0001, ***P<0.001, **P<0.01; two-tailed unpaired t-test, mean±SE. Representative images of DAPI-stained nuclei of additional strains are shown in Fig. S2A. B) Quantification of nuclear circularity index and nuclear abnormalities in three CAF strains infected with either a LacZ control (LacZ) or AR overexpressing lentivirus (AR OE). Cells were stained with DAPI and anti-AR antibody to determine nuclear morphology, and AR expression as shown in Fig. S2B and quantification was carried out as in Fig. 1B,C. For circularity, n=>50 cells per condition; for abnormalities, n=3 HDF strains; ****P<0.0001, **P<0.01; two-tailed unpaired t-test, mean±SE. C-D) Representative images and quantification of AR protein levels (C) and nuclear abnormalities (D) of three CAF strains treated with DMSO control (Ctrl) or Ostarine (10 µM) for 48 hours. Cells were stained with anti-AR antibodies and DAPI and quantifications were carried out as in the previous panels. n= 3 strains (>100 cells per condition), **P<0.01; two-tailed unpaired t-test. mean±SE. E) Immunofluorescence (IF) staining of excised skin SCC lesions with anti-pan-keratin (Red) and -Vimentin (red) antibodies and DAPI for cell type identification. The areas analyzed by 3D image reconstructions are indicated by a white dotted line, for SCC lesion, and solid line, for unaffected flanking skin. Parallel sections were stained with anti-vimentin and anti-AR antibodies for the analysis shown in (G) and with anti-vimentin, anti-lamin A/C and anti pSer301 - lamin A/C antibodies are shown in Fig. 5E. Scale bar 100 μm. F) Schematic representation of 3D reconstruction of Z-stack confocal images derived from the SCC versus distal skin areas indicated in the previous panel. G) 3D surface reconstruction images of DAPI-stained nuclei (cyan) of vimentin-positive fibroblasts (violet) concomitantly stained with anti-AR antibodies (red) from the SCC versus distal skin areas shown in (E). Scale bar: 5 μm. H-I) Quantification of (H) nuclear circularity index and (I) nuclear abnormalities in fibroblasts from SCC-associated versus distal skin, of three patient-derived skin SCCs. Quantifications are based on 3D image reconstructions and volume rendering of DAPI-stained nuclei using the Imaris software package. Images of patient #2 and 3 samples are shown in Fig. S2D,E,F. Quantification of levels of AR expression are shown in Fig. S2F. For circularity, n=>100 cells per condition; for abnormalities, n=3 HDF strains; ***P<0.001, **P<0.01 *P<0.05; two-tailed unpaired t-test, mean±SE.

To assess whether nuclear changes are also a feature of CAFs *in vivo*, we examined the nuclear shape of the fibroblasts within skin SCC lesions compared to fibroblasts from flanking unaffected skin in multiple patient-derived tissues. Double immunofluorescence analysis using antibodies against pan-keratin and vimentin was used to localize cancerous lesions and identify cell types (Fig. 2E; Fig. S2D). Parallel sections were stained with antibodies against vimentin and AR with DAPI for nuclear visualization. Confocal microscopy and 3D reconstruction of multiple optical planes showed that decreased AR expression in SCC-associated fibroblasts was accompanied by significant changes in nuclear shape (decreased circularity) and abnormalities (increased blebbing, micronuclei) (Fig. 2F-I, Fig. S2E,F).

Thus, nuclear changes are an AR-dependent feature of clinically occurring CAFs and occur both *in vitro* and *in vivo*.

### 3. AR is a determinant of nuclear lamin A/C localization

Lamin A/C plays a prominent role both at the level of the nuclear envelope and inside the nucleus, in control of gene transcription ^14, 30, 31^. Co-immunoprecipitation (Co-IP) and proximity ligation assays (PLAs) with antibodies against AR and lamin A/C showed an association of these proteins in HDFs, with positive PLA signals (puncta) being lost in cells with *AR* silencing (Fig. 3A,B, Fig. S3A). Number of PLA puncta was also drastically reduced in cells with degradation of the AR protein by a PROTAC-derived compound (ARCC4), while it was enhanced by treatment with the AR ligand dihydrotestosterone (DHT) that leads to increased AR expression (Fig. 3C, Fig. S3B).

**Figure 3.**
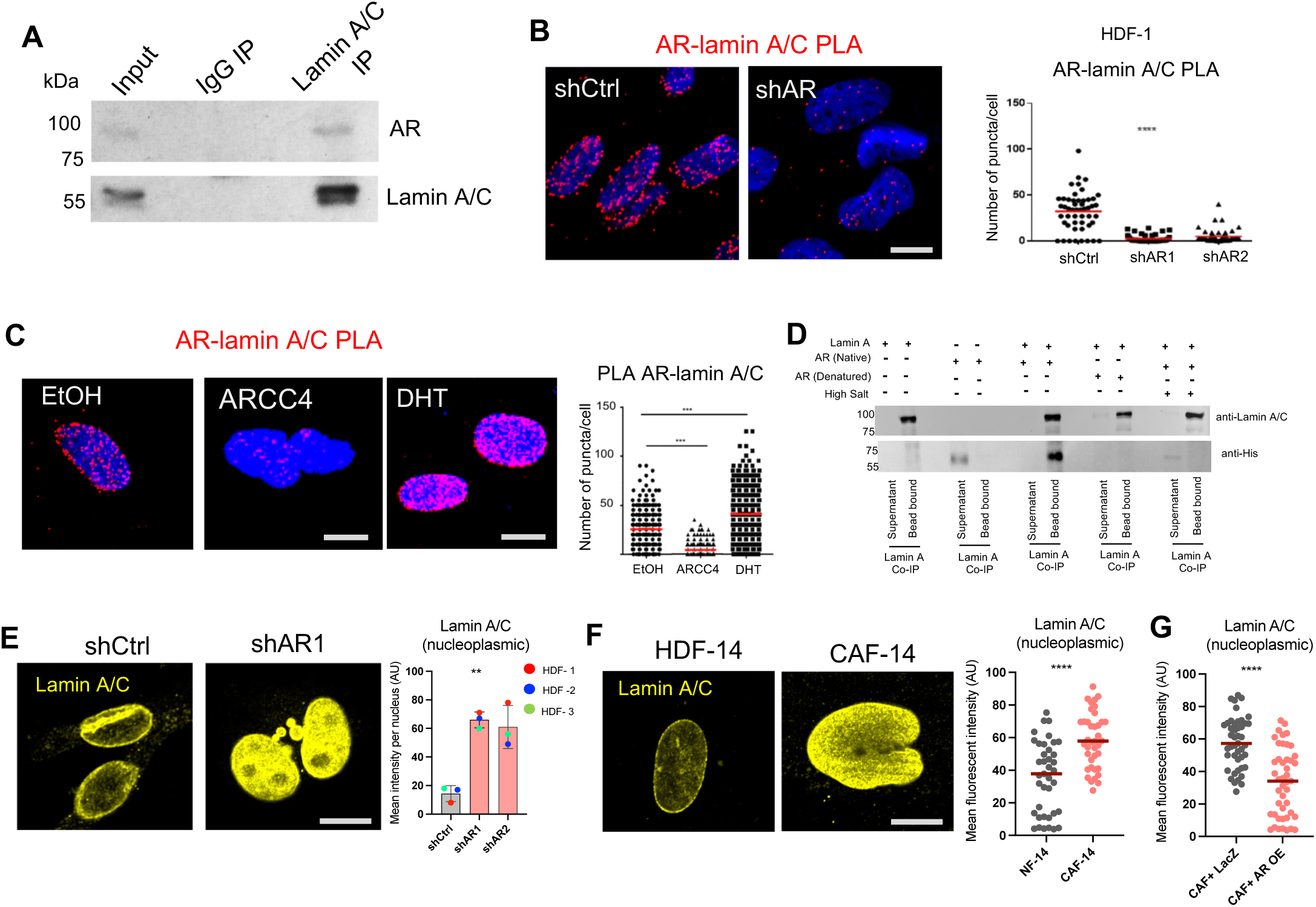
AR is a determinant of nuclear lamin A/C localization. A) Immunoprecipitation of HDF cell lysates with anti-lamin A/C antibodies versus nonimmune IgGs followed by SDS gel electrophoresis together with total lysate input and sequential immunoblotting with anti-AR and anti-lamin A/C antibodies. B) Proximity ligation assays (PLA) with antibodies against AR and lamin A/C in HDFs plus/minus shRNA-mediated *AR* silencing. Red fluorescence puncta resulting from the juxtaposition of anti-AR and lamin A/C antibodies were visualized by confocal microscopy with DAPI nuclear staining (blue). Shown are representative images and quantification of the number of puncta per cell as determined with ImageJ Analyze Particles tool. n(cells)>50 per condition, together with mean and statistical significance. ****p<0.001; non-parametric one-way ANOVA test. Scale bar: 5 µm. PLAs with two additional strains are provided in Fig. S3A. C) PLA with antibodies against AR and lamin A/C in HDFs treated with Ethanol vehicle alone, (EtOH), the AR degrading compound ARCC4 (1 µM) or AR agonist DHT (10 nM) for 72 h. Cells were cultured in medium containing charcoal stripped fetal bovine serum starting 24 h before the treatments. Shown are representative images and quantification of the number of puncta per cell, n(cells)>100 per condition together with mean and statistical significance. ****p<0.001, two-tailed unpaired t-test. Scale bar: 5 µM. Impact of ARCC4 and DHT treatment on levels of AR expression, as assessed by immunoblot analysis, is shown in Fig. S3B. D) *In vitro* AR-lamin A/C binding. Recombinant full-length lamin A protein (lamin A) was admixed with His-tagged AR protein (aa 1-560 aa) under native conditions. AR was previously denatured by a brief heat treatment (20 mins at 85^0^C), or proteins were incubated in high salt buffer (800 mM NaCl) as specificity controls. Immunoprecipitates with anti-lamin A/C antibody in parallel with 5% of the unbound supernatants were analyzed by immunoblotting with anti-lamin A/C or anti-His antibodies. E) Immunofluorescence analysis of lamin A/C localization in three HDF strains plus/minus *AR* silencing. Shown are representative images and quantification of nucleoplasmic lamin A/C fluorescence signal intensity assessed by Image J and expressed in arbitrary units (AU). n =3 strains (>50 cells per condition), **P<0.01; non-parametric one-way ANOVA test, mean ± SE. Scale bar 10 μm. F) Immunofluorescence analysis of lamin A/C localization in CAFs and matched HDFs from the same patient. Shown are representative images and quantification of nucleoplasmic lamin A/C fluorescence signal intensity assessed by Image J and expressed in arbitrary units (AU). n=>50 cells per condition together with mean and statistical significance, ****P<0.0001; two-tailed unpaired t-test, The data shown are the mean ± SE. Data for additional strains are shown in Fig. S3E. G) Quantification of nucleoplasmic lamin A/C in CAFs plus/minus AR overexpresson. CAFs were infected with either a LacZ control (LacZ) or AR overexpressing lentivirus (AR OE) lentivirus, selected and analyzed 7 days post infection by IF with anti lamin A/C antibodies. The nucleoplasmic distribution of lamin A/C was quantified as in Fig.3F. n=>50 cells per condition together with mean and statistical significance,****P<0.0001; two-tailed unpaired t-test.

To assess whether AR and lamin A/C can bind directly to each other in the absence of ligand, we incubated purified recombinant His-tagged AR protein, which lacks the ligand binding domain (UniPort: P10275-2; aa 1-560 amino acids), with full-length lamin A. Lamin A pull-down with anti-lamin A/C antibodies followed by immunoblotting showed effective binding of lamin A to the recombinant AR protein. AR/lamin A association was found to be lost when the AR protein was heat denatured prior to binding assay or when the binding reaction was performed in high salt (800 mM NaCl) buffer (Fig. 3D).

The AR-lamin A/C association is of functional significance. In fact, while total lamin A/C levels were unaffected by *AR* silencing (Fig. S3C), immunofluorescence analysis showed a marked accumulation of lamin A/C in the nuclear interior of multiple HDF strains with silenced *AR* (Fig. 3E), with no significant change in lamin B1 distribution (Fig. S3D). Nucleoplasmic accumulation of lamin A/C was also found by immunofluorescence analysis of CAFs compared to matched HDFs, which was reversed by overexpression of AR (Fig. 3F,G, Fig. S3E,F).

Thus, AR associates with lamin A/C and is a determinant of its proper nuclear distribution in both HDFs and CAFs.

### 4. AR loss compromises lamin A/C association with the PPP1 phosphatase and induces lamin A/C phosphorylation at Ser 301

Localisation and function of lamin A/C involves its association with a variety of proteins ^17, 32, 33^, which may be affected by loss of AR. We investigated this possibility by mass spectrometric analysis of lamin A/C immunoprecipitates from three different HDF strains plus/minus *AR* silencing. The association of lamin A/C with many proteins associated with the cytoskeleton, nuclear pore and importin complexes was consistently reduced by AR loss, together with enzymes involved in chromatin modification and RNA metabolism (Fig. 4A). Mass spectrometry also demonstrated a consistent decrease in the interaction between lamin A/C and three catalytic subunits of protein phosphatase 1 (PPP1C-A, B, C) as well as a regulatory subunit (PPP1R18) in HDFs with silenced *AR* (Fig. 4A). The AR-dependent association of lamin A/C with PPP1 was independently confirmed by co-immune precipitation and PLA assays with antibodies against lamin A/C and and PPP1C-A in HDFs with silenced *AR* versus controls (Fig. 4B,C, Fig. S4A). PLA assays showed also a marked reduction of lamin A/C - PPP1 puncta in HDFs treated with the AR-degrading ARCC4 compound versus ethanol control (Fig. 4D, S4B).

**Figure 4.**
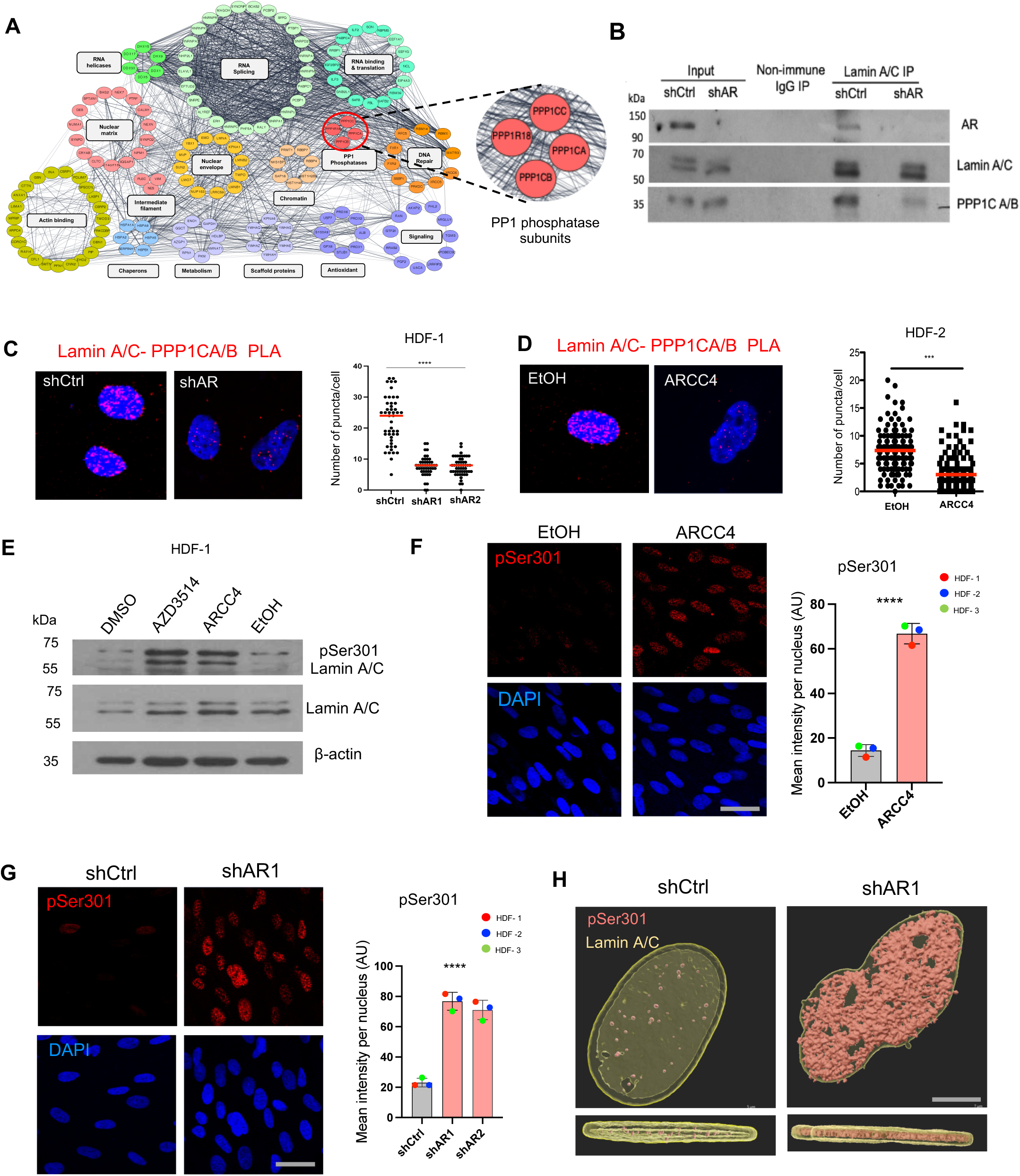
Loss of AR compromises the association of lamin A/C with the phosphatase PPP1 and results in increased lamin A/C phosphorylation at Ser 301. A) Cytoscape network visualization of proteins with consistently reduced association with lamin A/C in HDFs with silenced *AR*. Proteins were identified by Liquid Chromatography with tandem mass spectrometry (LC-MS/MS) analysis of lamin A/C immunoprecipitates from three different HDF strains plus/minus shRNA-mediated AR silencing. The spectral analysis identification of proteins, statistical validation, and scoring were carried out using the Mascot search engine (www.matrixscience.com). Shown are the names of lamin A/C associated proteins with reduced association (log fold change >1.5) in at least two of the three HDF strains with silenced AR, with functional classification carried out using Cytoscape 3.8.2. The inset shows the PPP1 subunits that were found to be associated with lamin A/C in HDFs in an AR dependent manner. B) Co-Immunoprecipitation assays with HDFs plus/minus shRNA-mediated *AR* silencing based on immunoprecipitation with anti-lamin A/C antibodies or nonimmune IgGs followed by sequential immunoblotting with anti-AR, -lamin A/C, and - PPP1CA/B antibodies, with parallel analysis of the inputs. C) Proximity ligation assays (PLA) with anti-lamin A/C and PPP1CA/B antibodies of HDFs plus/minus shRNA-mediated *AR* silencing. Shown are representative images and quantification of the number of puncta per cell. n(cells)>50 per together with mean and statistical significance, ****p<0.0001, non-parametric one-way ANOVA Kruskal–Wallis test. Scale bar: 10 µm. D) PLA with anti-lamin A/C and PPP1CA/B antibodies of HDFs treated with the AR degrading compound ARCC4 (1 µM) or solvent DMSO control for 72 h. Shown are representative images and quantification of the number of puncta per cell. n(cells)>50 per together with mean and statistical significance, ***p<0.001, unpaired student t-test. Scale bar: 10 µm. E) Immunoblot analysis of HDFs treated with the AR inhibitors ARCC4 (1 µM) or AZD3514 (10 uM) in parallel with indicated solvent controls for 72 h, with antibodies against phospho-Ser301 laminA/C, total lamin A/C and β-Actin as equal loading control. F) Immunofluorescence analysis of three HDF strains treated with ARCC4 (1 µM) or ethanol vehicle alone (EtOH) for 72 h with antibodies against phospho-Ser301-lamin A/C (pSer301; red) with DAPI for nuclear staining. Shown are representative images and quantification of phospho-ser301-lamin A/C fluorescence signal intensity assessed by Image J and expressed in arbitrary units (AU). n= 3 strains (>50 cells per condition), ****p<0.0001, unpaired student t-test, mean ± SE. Scale bar: 20 µm. G) Immunofluorescence analysis of three HDF strains infected with two different *AR* silencing lentiviruses versus virus control with antibodies against phospho-Ser301-lamin A/C (pSer301; red) and DAPI for nuclear staining. Shown are representative images and quantification of phospho-ser301-lamin A/C fluorescence signal intensity assessed by Image J and expressed in arbitrary units (AU). n= 3 strains (>50 cells per condition), ****p<0.0001, non-parametric one-way ANOVA Kruskal–Wallis test, mean ± SE. Scale bar: 20 µm. Fluorescence intensities per individual counted cells for each strain are shown in Fig. S4D. H) 3D surface reconstruction of whole nuclei of HDFs plus/minus *AR* silencing stained with anti-phospho-S301-lamin A/C (red) and total lamin A/C (yellow) antibodies. Z-stacks of confocal images were used for 3D volumetric surface reconstruction using the Imaris Surface tool. Shown are top and lateral views of the 3D reconstructions. Scale bar: 5 µm.

PPP1 is involved in control of lamin A/C phosphorylation ^34, 35^, which is in turn a key determinant of its localisation and function ^36^. A number of phosphorylated sites were detected in the mass spectra profiles of lamin A/C immunoprecipates, among which phosphorylation at Ser 301 was consistently enriched in all three HDF strains with silenced *AR* compared to control (Fig. S3C). Immunoblot analysis with phospho-specific antibodies showed a marked increase in lamin A/C phosphorylation at Ser 301 in HDFs treated with the AR inhibitors AZD3514 and ARCC4 (Fig. 4E). The induction of Ser301 phosphorylation was confirmed by immunofluorescence analysis of several HDF strains treated with the ARCC4 compound in parallel with the PPP1 inhibitor tautomycetin ^37^ (Fig. 4F, Fig. S4D), and was similarly observed upon *AR* silencing in multiple HDF strains (Fig. 4G, Fig. S4E). 3D surface reconstruction of confocal Z-stack images showed that phospho-Ser301-lamin A/C induced by *AR* silencing localizes to the nuclear interior, while total lamin A/C is also present at the membrane (Fig. 4H).

Thus, loss of AR in HDFs impairs lamin A/C association with the PPP1 phosphatase and results in increased lamin A/C phosphorylation at Ser301.

### 5. Lamin A/C phosphorylation at Ser301 is a feature of CAFs

An important question was whether the observed increase(s) in lamin A/C phosphorylation at Ser301 also occurs in CAFs. Immunofluorescence analysis using phospho specific antibodies revealed a substantial elevation of Ser301 phosphorylation, with pronounced nucleoplasm distribution, in several CAF strains compared to matched HDFs from the same patients (Fig. 5A, Fig. S5A). In parallel with the restored nuclear shape, Ser301 phosphorylation was significantly reduced by treatment of CAFs with the AR agonist ostarine (Fig. 5B), with a concomitant increase in PPP1-lamin A/C association (Fig. S5B).

**Figure 5.**
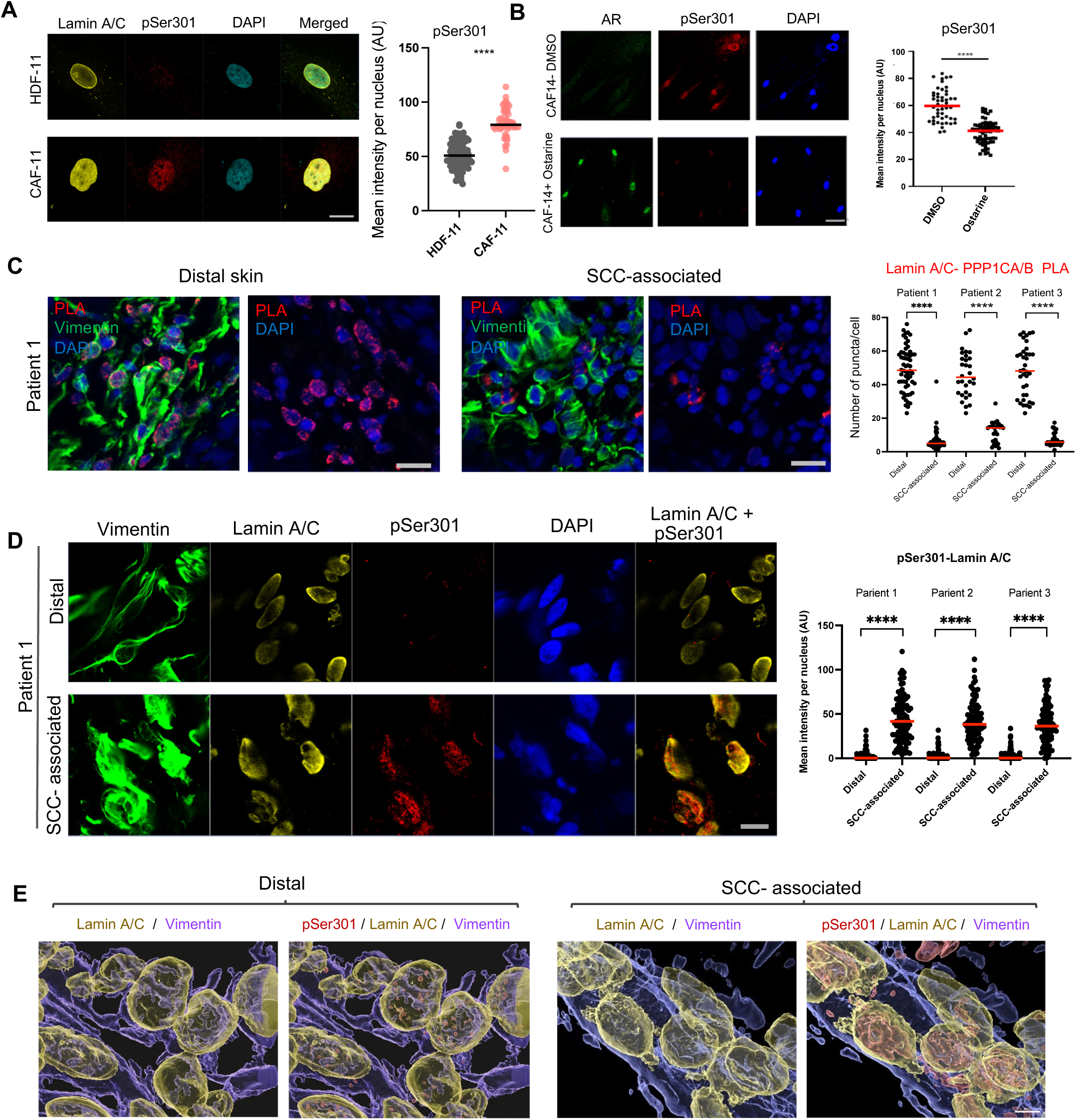
Lamin A/C phosphorylation at Ser 301 is a signature of CAFs. A) IF analysis of CAFs and matched normal fibroblasts (HDFs) from the same patient with images of pSer301 lamin A/C (pSer301; red) and total lamin A/C (yellow). Shown are representative images and quantification of phospho-ser301-lamin A/C fluorescence signal intensity assessed by Image J and expressed in arbitrary units (AU). n(cells)>50 per condition together with mean and statistical significance, ****p<0.0001, unpaired student t-test. Scale bar: 10 µm. Similar analysis of two different CAF strains and matched HDFs is shown in Fig. S5A. B) Double IF analysis of CAFs, treated with the AR agonist Ostarine (10 μM) or DMSO vehicle alone for 48 h, with antibodies against AR (green) or phospho-Ser310 lamin A/C (red). Shown are representative images and quantification of phospho-ser301-lamin A/C fluorescence signal intensity assessed by Image J and expressed in arbitrary units (AU). n(cells)>50 per condition together with mean and statistical significance,****p<0.0001, unpaired student t-test. Scale bar: 20 µm. C) PLA of skin SCC-associated fibroblasts versus fibroblasts in distant unaffected skin with antibodies against lamin A/C and PPP1CA/B (red puncta), with concomitant IF with anti-vimentin antibodies for cell type identification (green). Shown are representative images and quantification of PLA signal in vimentin positive cells in matched pairs of skin SCC associated versus distal skin fibroblasts from three different patients. n(cells)>20 per together with mean and statistical significance, ****p<0.0001, unpaired student t-test. Scale bar is 30 µm. D) IF analysis of skin SCC-associated fibroblasts versus fibroblasts in distal unaffected skin with antibodies against total (yellow) and phospho-Ser301 (red) lamin A/C, with anti-vimentin antibodies for cell type identification (green). Shown are representative images and quantification of anti-phosphoSer301 signals in matched pairs of skin SCC-associated versus distal skin fibroblasts from three different patients (parallel sections from those analyzed in the previous panel) n= cells >50 per condition together with mean and statistical significance, ****p<0.000, paired t-test. Scale bar: 30 µm. E) 3D volumetric surface reconstruction of confocal IF images of the SCC lesion and unaffected flanking skin as shown in Fig. 2E with antibodies against phospho-S301 (red) and total lamin A/C (yellow) and vimentin (purple), using the Imaris Surface tool, Manual mode. For each individual sample, threshold parameters were tested and set based on the weakest positive signal. Scale bar: 5 µm.

The results also apply to the *in vivo* situation. Tissue PLA demonstrated a significant reduction in lamin-PPP1CA/B association in lesion fibroblasts in skin SCCs compared to fibroblasts in distal unaffected skin from the same patients (Fig. 5C). Parallel, IF analysis revealed a significant increase in Ser301 phosphorylation and nucleoplasm distribution in the SCC-associated fibroblasts versus fibroblasts from distal skin (Fig. 5D, Fig. S5C). Confocal microscopy followed by 3D reconstruction confirmed these findings, showing a pronounced intranuclear distribution of phospho-Ser301 lamin A/C in SCC-associated fibroblasts (Fig. 5E).

Thus, increased lamin A/C phosphorylation at Ser301 is a feature of CAFs *in vitro* and *in vivo*, which is accompanied by impaired lamin A/C – PPP1 association.

### 6. AR loss results in enhanced binding of phospho-Ser 301 lamin A/C to CAF effector genes in concomitance with chromatin activation

Polymeric lamin A/C is known to interact with large heterochromatin domains called lamina-associated domains (LADs) ^38^, while dimeric nucleoplasmic lamin A/C can bind to a number of genomic sites characteristic of active enhancers ^39^. To assess to which extent AR loss leads changes in chromatin configuration that are associated with lamin A/C redistribution, HDFs plus/minus *AR* silencing were analyzed by ChIP-seq with antibodies against total and phospho-Ser301 lamin AC in parallel with antibodies against H3K9me3 and H3K27ac as markers of closed versus open chromatin configuration, respectively ^40^. ATAC-Seq (assay for transposase accessible chromatin with sequencing ^41^ was also employed to assess AR-dependent changes in chromatin accessibility across the genome.

Consistent with the observed increase in lamin A/C nucleoplasmic localization and Ser301 phosphorylation, ChIP-seq analysis revealed a marked global increase in narrow peaks of total and phospho-Ser301 lamin A/C in *AR* silenced HDFs (Figure 6A).

**Figure 6.**
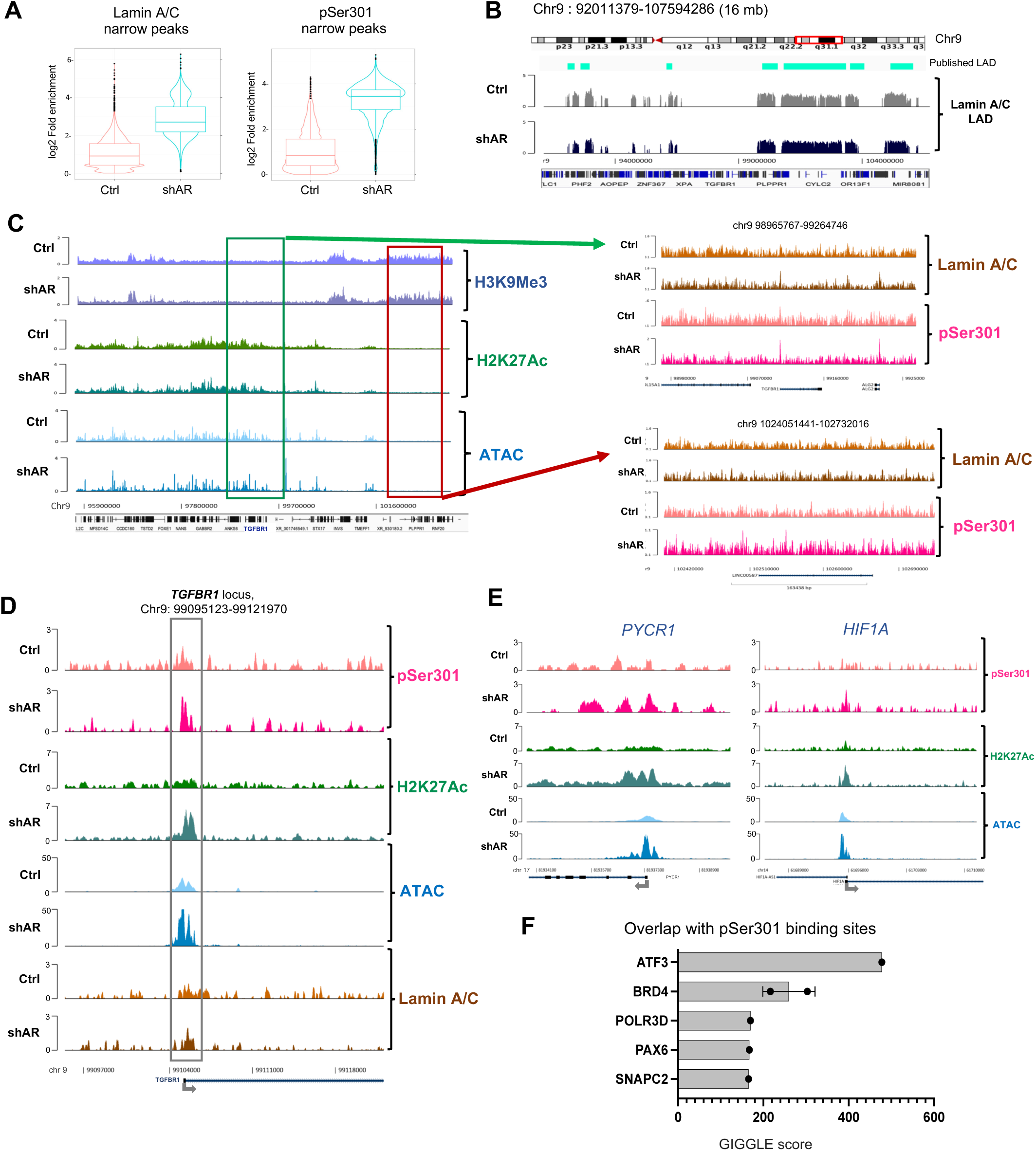
AR is a determinant of chromatin association of total and phospho-Ser 301 lamin A/C. A) Violin plots showing the enrichment of narrow peaks of total and phospho-Ser301 binding to chromatin, as assessed by ChIP-seq analysis of two different HDF strains with shRNA-mediated *AR* silencing versus control. B) Genomic tracks of lamin-associated domains (LAD) of a representative chromosomal region (16 Mb, chromosome 9 q31.1, red box), detected by ChIP-seq analysis with anti-lamin A/C antiboides of HDFs plus/minus *AR* gene silencing after high stringency chromatin extraction conditions. Position of experimentally determined LADs was calculated using Enriched Domain Detector (EDD) software 42 and aligned with that of published LADs from IMR90 fibroblasts (Guelen et al., 2008), depicted in cyan bars on top. Lower bar: indication of genes or transcribed regions (black lines) lying within the examined region. C) Parallel analysis of chromatin configuration and total and phospho-lamin A/C binding in HDFs plus/minus *AR* gene silencing. <u>Left</u> : ChIP-seq profiles of H3K9Me3 and H3K27Ac binding, as markers of heterochromatin and euchromatin respectively, and ATAC-seq profiles, to determine regions of chromatin accessibility, of a genomic region (7 mb) of functional significance, encompassing the *TGFBR1* gene as indicated in red in the lower bar together with other genes . <u>Right</u> : ChIP-seq profiles of total and phospho-lamin A/C binding to the open and closed chromatin areas marked in the left panel as green and red squares, respectively. D) Higher resolution analysis of the *TGFBR1* gene and flanking region. ChIP-seq and ATAC-seq profiles were uploaded onto the EaSeq tool (https://easeq.net/) with the ENCODE database for human fibroblasts as reference, to map binding peaks on 30 KB genomic region encompassing the *TGFBR1* gene. The boxed area corresponds to the 1500 bp promoter region of the gene around the Transcription Start Site (TSS). E) Similar high-resolution analysis as in the previous panel of two other CAF effector genes of relevance, *PYCR1* and HIF-1, as discussed in the text. Position of the TSS and transcribed region is indicated. Analysis of other genes of interest is provided in Fig. S6. F) Prediction of transcription factors binding to chromatin in concomitance with phospho-Ser 301 lamin A/C in human fibroblasts. GIGGLE score provides an overlap analysis between user-defined peak profiles and deposited ChIP-seq profiles in the Cistrome DB data base (http://dbtoolkit.cistrome.org/). Shown are the transcription factors and epigenetic regulators with top ranking GIGGLE scores overlapping with phospho-Ser301 Lamin A/C binding peaks in ChIP-seq profiles of human fibroblasts. Dots correspond to individual data set.

LAD formation was investigated by Enriched Domain Detector (EDD) analysis ^42^ of lamin A/C ChIP-seq profiles of HDFs following high stringency nuclear extract conditions to improve detection of stronger DNA-lamin association ^43^. As shown in Fig. 6B, the experimentally determined distribution of LADs in HDFs was matching that of previously reported LADs of IMR90 fibroblasts ^42^, with only minimal changes occurring in smaller LADs in HDFs with silenced *AR*.

To analyse changes in chromatin modifications, ChIP-seq profiles obtained from HDFs following standard nuclear extraction conditions were uploaded to the Easeq tool, an interactive software for analysis and visualization of ChIP-sequencing data (https://easeq.net/). Heterochromatin regions with elevated H3K9me3 binding showed a similar distribution in HDFs with silenced *AR* as in controls (Fig. 6C, left). The overall localisation of euchromatin regions marked by elevated H3K27ac and ATAC-seq peaks was also maintained, with changes in peak intensity, indicative of more specific variations in chromatin accessibility (Fig. 6C, left). Importantly, *AR* silencing resulted in a large number of new peaks of total and phospho-S301 lamin A/C binding in both hetero- and euchromatin regions (Fig. 6C, right).

The functional significance of increases in lamin A/C binding to heterochromatin regions will have to be separately investigated. Here we focused on the changes in lamin A/C binding to euchromatin regions and more specifically to loci of CAF effector genes induced by *AR* silencing. As a point in case, we examined genomic regions encompassing the *TGFBR1* gene, whose expression and function are enhanced in CAFs ^4, 44^. *AR* silencing resulted in a notable increase of phospho-Ser301 lamin A/C binding peaks at a 1.5 kb region encompassing the transcription start site (TSS) of the gene, with a striking co-occurrence of enhanced H3K27ac binding and chromatin accessibility (ATAC peaks) (Fig. 6D). There were also changes in total lamin A/C binding peaks, less clustered at the *TGFBR1* promoter region and coinciding to a limited extent with those of phospho-Ser301 (Fig. 6D).

A similar coincidence of H3K27ac, ATAC and phospho-lamin A/C binding peaks was found at the promoter regions of many other CAF effector genes under AR control, including *PYCR1* (Pyrroline-5-Carboxylate Reductase 1), a key enzyme for proline synthesis recently linked to CAF metabolic activation ^45^, *HIF-1α* (hypoxia-inducible factor-1α), a transcription factor important for CAF activation in aggressive tumors ^46^ and genes with matrix remodeling function (*PLOD2*, *COL1A1*, *COL5A1*, *FBN1*) or coding for key growth factors and immune modulators (*FGF2*, *IL11*, *CXCL5*, *INHBB*) ^47^ (Fig. 6E, Fig. S6).

For further mechanistic insights, we performed an unbiased analysis of publicly available ChIP-seq profiles of known transcription factors and chromatin regulators in human fibroblasts using the Cistrome Data Browser and tool kit (http://dbtoolkit.cistrome.org/) ^48^. The analysis revealed a highly significant overlap of phospho-Ser301 lamin A/C binding peaks in AR-silenced HDFs with those of a restricted number of proteins with a known role in transcription, including ATF3 and BRD4, two known regulators of CAF activation (Kim et al., 2017) (Fig. 6F).

Thus, *AR* silencing in HDFs results in changes in chromatin activation associated with enhanced phospho-Ser301 lamin A/C binding of likely significance for CAF activation.

### 7. Expression of a phosphomimetic Ser 301 lamin A/C mutant is sufficient to trigger CAF activation unlinked from stromal fibroblasts senescence

To probe into the specific consequences of lamin A/C Ser 301 phosphorylation, we expressed a phosphomimetic Ser 301 to Asp (S301D) mutant and wild-type (WT) lamin A/C via lentiviral vector transduction. Infection of multiple HDF strains with lentiviruses expressing the S301D mutant or wild-type lamin A/C had no effects on proliferation and did not induce the senescence-associated changes in cell morphology that are observed upon *AR* gene silencing ^21^ (Fig. S7A).

Global transcriptomic analysis showed consistent changes in gene expression with the upregulation of many CAF effectors in two different HDF strains expressing the S301D mutant versus wild-type lamin A/C (Fig. 7A). RT-qPCR analysis confirmed that expression of the S301D lamin A/C mutant in HDFs induces multiple CAF effector genes without up-regulation of *CDKN1A*, a determinant of senescence induced by *AR* silencing in these cells ^21^ (Fig. S7B). More general, gene set enrichment analysis (GSEA) revealed that profiles of HDFs expressing the S301D mutant were significantly enriched for gene signatures related to CAF activation of the matrisome and myofibroblast types ^7, 49^ (Fig. 7B). A signature of up-regulated genes in HDFs with S301D lamin A/C mutant expression was established and used to analyze recently reported single-cell RNA-seq profiles of head and neck squamous cell carcinoma (HNSCC)^50^. A high signature score was found specifically in the CAF and myofibroblast subpopulations (Fig. 7C).

**Figure 7.**
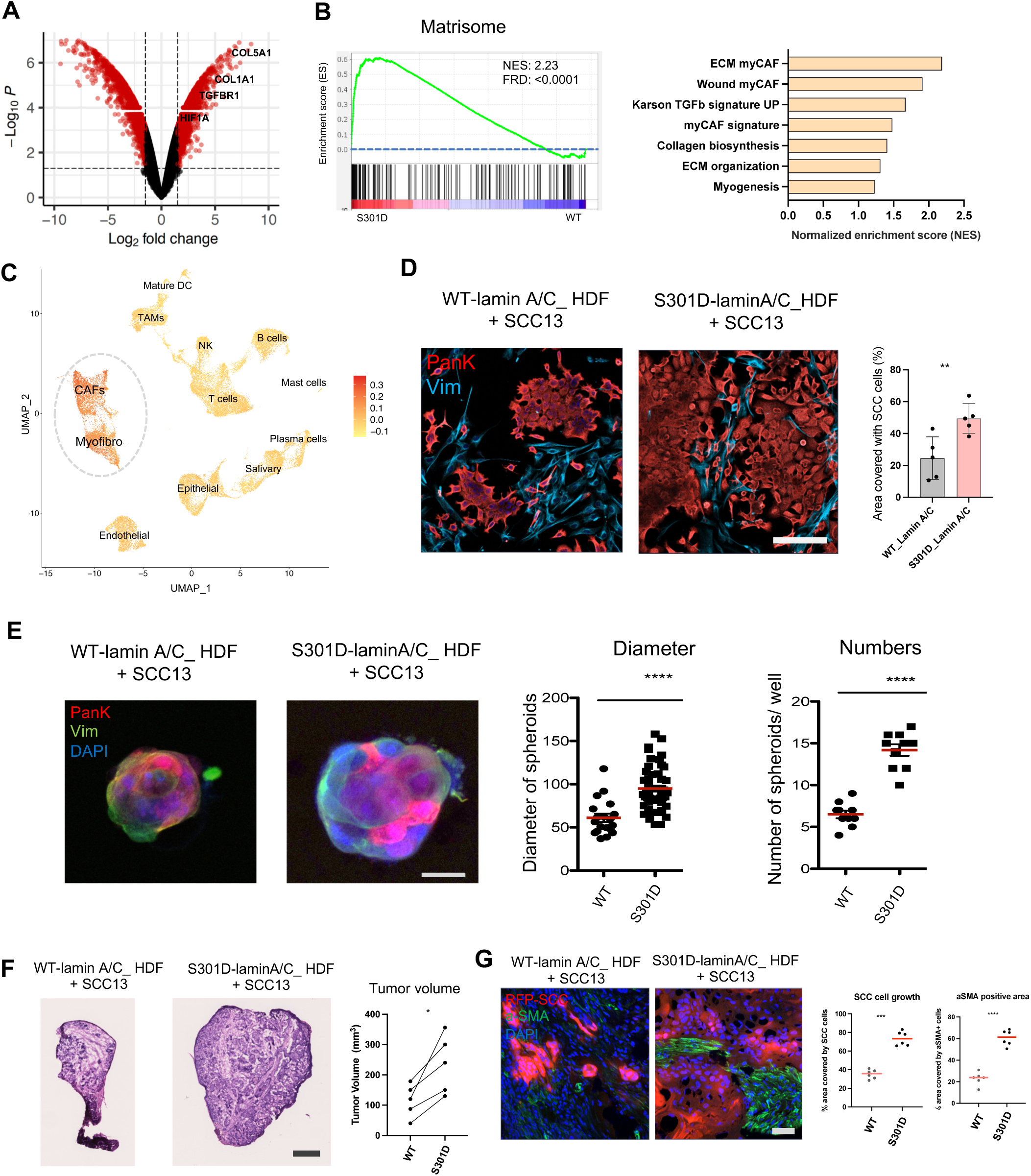
Expression of a phosphomimetic Ser 301 lamin A/C mutant is sufficient to trigger CAF activation. A) Volcano-plot of differentially expressed genes in HDFs expressing a phosphomimetic Ser to Asp 301 mutant (S301D) versus wild-type lamin A/C. Two HDFs strains were stably infected with lentiviruses expressing a S301D mutant versus wildtype lamin A/C (WT). Eight days post infection transcriptomic analysis was carried out with Clariom D cDNA array hybridization. Differentially expressed genes (fold changes >1.5; p values <0.05) are highlighted in red. The names of a few upregulated genes by expression of the S301D lamin A/C mutant are indicated. B) Gene set enrichment analysis (GSEA) of transcriptomic profiles of HDFs expressing the S301D mutant versus wild type lamin A/C. Left; shown is the positive enrichment plots of the CAF matrisome gene signature (GSE71340)). The position of each gene is indicated by black vertical bars; the enrichment plot is shown in green, together with normalized enrichment score (NES) and FRD-q values. Right; bar plot indicates additional gensets of CAF-related pathways enriched in transcriptomic profiles of HDFs expressing the S301D mutant. Signatures were downloaded from the GSEA web page (http://www.gsea-msigdb.org/gsea/msigdb/human/genesets.jsp), and the TGFb signature (GSE79621), CAF gene signature (GSE122372) and my CAF (GSE93313) was downloaded from respective GEO. C) Enrichment of the S301D lamin A/C gene signature on CAF and myofibroblast populations of head and neck squamous cell carcinoma (HNSCC) single cell clusters. Shown is the Uniform Manifold Approximation and Projection (UMAP) of different cell type clusters of HNSCC tumours ^50^ overlaid with the AddModule score of the top 200 upregulated genes in S301D expressing cells. Each dot represents a single cell and colour corresponds to the relative expression of the S301D signature within a cell. Yellow corresponds to little/no gene expression, while red corresponds to increased gene expression as indicated in the scale. D) Cancer cells expansion assays. HDFs infected with lentiviruses expressing the S301D mutant versus wild-type lamin A/C were co-cultured with SCC13 cells on a thin of matrigel layer for 4 days. Immunofluorescence analysis with anti-Pan-keratin (red) and -Vimentin (blue) antibodies was used for cell type identification. Percentage of SCC positive area was quantified by Image J analysis of 5 random fields per well. n=5 independent replicate wells. Mean ± SD; unpaired t-test. ***p<0.001, Scale bar: 100 µm. E) Spheroid formation assays. HDFs expressing the S301D mutant versus lamin A/C were co-cultured in 3D with SCC13 cells in 10% Matrigel-coated 8 well glass slide for 7 days. IF with anti-Pan-keratin (red) and -Vimentin (blue) antibodies were used for cell type identification. Shown are representative images and quantification of diameter and number of spheroids by ImageJ analysis of digitally acquired images. For diameter, n=>25 spheroids per condition; for spheroid numbers, n=8 wells. Mean ± SD, unpaired t-test.*** p<0.001, Scale bar: 500 µm. F) *In vivo* tumor formation assays. RFP-expressing SCC13 cells were co-injected with HDFs expressing S301D mutant versus wild-type lamin A/C intradermally into contralateral mouse back skin. Mice were sacrificed 14 days later. Shown are representative images of H&E stained lesions and quantification of excised tumor volumes calculated using the formula V = 3.14 × (W2 × L)/6 V= volume, W= width, and L= length. n=5 mice. Scale bar is 500 µm. G) IF analysis of RFP-expressing SCC cells and a-SMA positive fibroblasts respectively. Shown are representative images and quantification of RFP and a-SMA-positive area per lesion, by ImageJ analysis of digitally acquired images. n=5 mice. Mean ± SD, unpaired two-tailed t-test, ***, p < 0.005; ****, p<0.0001, Scale bar is 100 µm.

A key property of CAFs is to enhance proliferation and tumorigenicity of neighboring cancer cells. In co-culture assays in thin Matrigel layers ^21^, expansion of squamous cell carcinoma (SCC13) cells was enhanced in the presence of HDFs with expression of the S301D mutant versus wild-type lamin A/C (Fig. 7D), with similar enhancing properties occurring in 3D spheroid forming assays (Fig. 7E). *In vivo* tumor-promoting activity was assessed by an orthotopic model of skin cancer formation based on intradermal injections of cells in Matrigel in the back skin of mice. As shown in Fig. 7F and Fig. S7C, SCC13 cells in combination with HDFs expressing the S301D mutant formed tumors of significantly larger size than with HDFs expressing wild-type lamin A/C. Lesions formed in the presence of S301D – expressing HDFs were characterized by greater density of SCC cells (expressing the RFP marker) with an increase of α−SMA positive fibroblasts (Fig. 7F,G).

Thus, expression of a phospho-mimetic Ser 301 lamin A/C mutant in HDFs is sufficient to induce a transcriptional program of CAF activation and confer on cells growth- and tumor enhancing properties.

## DISCUSSION

CAFs are key determinants of the cancer process, from initiation to metastatic spread ^6, 51^. Resident stromal fibroblasts are a primary source of CAFs, with a plastic response to various cues resulting in CAF subpopulations with distinct functions ^4–9^. The androgen receptor (AR) represses a senescence program of gene transcription associated with early steps of CAF activation and acts as a master negative regulator of CAF effector genes ^21^. Here we have uncovered a novel role of AR as a determinant of nuclear envelope integrity, lamin A/C distribution, phosphorylation, and transcription modulatory function of relevance for CAF activation.

An involvement of AR in the maintenance of nuclear structure, to our knowledge, has not been previously reported. We have shown that AR loss in normal dermal fibroblasts by either genetic or pharmacologic approaches results in significant deformation of nuclear shape, nuclear abnormalities (blebbing, micronuclei formation) and ruptures during interphase. These are also a feature of CAFs, both in culture and in cancer lesions, which can be counteracted by increased AR expression. By analogy with what has been proposed for cancer cells ^2, 52, 53^, nuclear abnormalities in CAFs may contribute to alterations in gene expression and genomic instability as they have been experimentally observed ^12, 13, 54, 55^.

Molecularly, the nuclear alterations resulting from decreased AR levels or loss can be explained by the fact that AR binds to nuclear lamin A/C and ensures its proper localization and association with proteins involved in nuclear membrane composition and connection with the cytoplasm. Mass Spec analysis of multiple HDF strains revealed that the association of many of these proteins with lamin A/C was consistently decreased upon *AR* silencing. For further biochemical and functional studies, we focused on the AR-dependent association of lamin A/C with the PPP1 phosphatase, given its potentially broader involvement in the modulation of lamin A/C function ^34, 35^.

Phosphorylation of nuclear lamins controls multiple aspects of their function, including depolymerization at the onset of mitosis, subnuclear distribution during interphase, chromatin organization, and gene transcription ^30^. The impact that changes in lamin A/C phosphorylation have on CAF activation, to our knowledge, has not been explored. A number of different kinases including PKC, MAPK, and Akt have been shown to phosphorylate lamins at different sites, leading to different outcomes that are better characterized in relation to the cell cycle ^30^. The PPP1 and PPP2 phosphatases have been implicated in the dephosphorylation of lamins, specifically in post-mitotic cells ^56^.

PPP1 is composed of multiple catalytic (PPP1CA, PPP1CB and PPP1CC) and regulatory subunits implicated in target specificity ^57^. In our experimental setting, we detected association of lamin A/C with multiple PPP1 subunits and found it to be significantly compromised by *AR* gene silencing or PROTAC-mediated AR degradation. Concomitantly, loss of AR resulted in a marked increase of lamin A/C phosphorylation at Ser 301, which was also observed upon PPP1 inhibition. Phosphorylation of lamin A/C at another residue implicated in the cell cycle, Ser22 ^39^, was observed in HDFs upon *AR* gene silencing but not PROTAC-mediated degradation, indicating that it is a more indirect consequence of AR loss (Fig. S4F,G). Increased phosphorylation at Ser301, but not Ser22, together with reduced lamin A/C - PPP1 association, was also observed in CAFs, in culture and in clinically occurring lesions, *in vivo*. The significance of lamin A/C phosphorylation at Ser 301, as reported in one study by the Akt kinase ^58^, has been explored to a very limited extent. By the combination of biochemical and direct functional approaches discussed below, we have shown that this modification is of importance for the transcriptional program of CAF activation.

Nuclear lamins have been extensively studied for their contribution to the control of gene expression through both direct and indirect mechanisms. Lamin-associated domains (LADs) are global determinants of chromatin organization. LADs are categorized into large chromatin regions that associate with the nuclear lamina in all cell types (constitutive, cLADs) and others that associate in a cell type-specific manner (facultative, fLADs) ^59, 60^. The distribution of LADs determined by lamin A/C ChIP-seq analysis of HDFs was only minimally affected by *AR* silencing. There was increased binding of total and phospho-Ser301 lamin A/C to both hetero- and euchromatin regions without changes in their overall localization. Focusing on euchromatin changes of more direct relevance for CAF activation, we found a striking co occurrence of increased pSer301 lamin A/C binding and peaks of open chromatin configuration and accessibility induced by AR silencing at the transcriptional promoter regulatory regions of the *TGFBR1* gene and multiple other CAF effector genes. The findings are of likely functional significance as the expression of a phosphomimetic Ser/Asp 301 mutant resulted in expression of these genes as well as a wider transcriptional program of CAF activation related to the myofibroblast subtype ^9, 61^. Strikingly, expression of the phosphomimetic mutant was by itself sufficient to confer upon HDFs the ability to enhance proliferation and tumorigenicity of neighboring cancer cells.

Previous studies on deregulated gene transcription resulting from nuclear lamin alterations in pathologic conditions implicated global changes in chromatin organization and looping as possible underlying mechanisms ^3, 14^. In the case of CAF activation, we propose that the center stage is taken by more selective changes at the level of promoter regions of myofibroblast CAF effector genes, whereby phospho-Ser301 lamin A/C binding results in their increased expression. The close overlap with binding peaks of ATF3 and BRD4, two key regulators of CAF gene transcription (Kim et al., 2017) point to an exciting interconnection to be explored in further studies in the context of actionable targets for stroma-focused anti cancer intervention.

## ACKNOWLEDGMENTS

We thank Tatiana Proust for technical assistance during the experiments, Paola Ostano and Markus Kirolos Youssef for bioinformatic analysis of the LADs, An Buckinx for critical reading of the manuscript, Anastasia Samarkina and other members of the laboratory for insightful discussions and suggestions. The Central Imaging Facility (CIF), the Electron Microscopy Facility (EMF) and the Protein Analysis Facility (PAF) of the University of Lausanne are gratefully acknowledged for their services and support. This study was supported by grants from the Swiss National Science Foundation (310030B_176404 “Genomic instability and evolution in cancer stromal cells„) and the NIH (R01AR039190, the contents do not necessarily represent the official views of the NIH). J.I. is supported by the European Union’s Horizon 2020 Research and Innovation Programme under Marie Skłodowska-Curie Grant Agreement No 859860.

## Authors contributions

SG, JI, LM, AT and CS performed experiments and/or analyzed the results with GPD. SG and GPD designed the study and wrote the manuscript.

## Methods

### Cell culture

HDFs, CAFs and SCC13 cells were maintained in Dulbecco’s modified Eagle’s medium (DMEM; Gibco, Cat# 31966021) supplemented with 10% fetal bovine serum (FBS; Gibco, Cat# 10270106) and 1% Penicillin-Streptomycin (BioConcept; Cat# 401F00H). Cells were cultured at 37°C and 5% CO2, and routine mycoplasma testing was performed. All experiments were conducted using cells from passages 4 to 10.

HDFs were prepared from the foreskin of healthy males (aged 1-5 years) with institutional approvals and informed consent (UNIL; protocol # 222-12), as previously described ^22^. Briefly, 10 mg/mL Dispase (Merck, Cat# D4693-1G) was used to digest the tissue and separate the dermis from the epidermis. The epidermis was then digested using 1% collagenase (Merck, Cat# C9891) for 1 hour at 37°C, followed by addition of the tissue extract was filtered through a 70 µm sieve, centrifuged, resuspended in complete medium and plated at a density of 1×10^5^ cells/10 cm dish culture. The primary HDF strains used in this study, AK1, PB2 and AK2, were designated as HDF-1, HDF-2 and HDF-3, respectively. Please refer to the Table S1 for a comprehensive list of all primary cells and cell lines used in this study.

Cancer-associated fibroblasts (CAFs) and matched normal fibroblasts (HDFs) were isolated from discarded skin tissue samples of squamous cell carcinoma (SCC) and flanking unaffected areas, respectively, obtained from the same (anonymized) patients with institutional approval (2000P002418) at Massachusetts General Hospital (Boston, MA, USA). Tissue sections were cut into small pieces (1-2 mm) and incubated with 0.25 mg/ml Liberase TL (Roche, Cat# 05401020001) at 37°C for 40 minutes. After administration of FBS to stop the reaction, the tissues were filtered through a 70 µm sieve, and centrifuged at 300 g.

Skin SCC13 tumour cells were originally reported in ^62^. For *in vivo* tumourigenesis assays, SCC13 cells were stably infected with an RFP-expressing lentivirus as previously described ^63^.

To culture cells on a soft matrix, HDFs were seeded onto collagen hydrogel-coated coverslips (Softslip; Cell Guidance Systems Ltd; Cat # SS24-COL-25) and allowed to grow for 48 hours under standard culture conditions.

### Cell Manipulations

#### Short hairpin RNA (shRNA)-mediated silencing of androgen receptor (AR)

Lentiviral particle production and *AR* silencing was performed following the previously described protocol ^21^. Briefly, HDFs were transduced with lentiviruses expressing *AR* gene silencing shRNAs (shAR1 (TRCN000000003718) 5’-CACCAATGTCAACTCCAGGAT-3’; shAR2 (TRCN000000003715): 5’-CCTGCTAATCAAGTCACACAT-3’, Sigma-Aldrich) in parallel with an empty pLko1 control vector (shCtrl; Sigma-Aldrich, Cat # NCLMIR001). The transduction was performed in DMEM medium supplemented with Polybrene (0.8 µg/ml) for a duration of six hours. Two days after infection, HDFs were selected for puromycin resistance (1 µg/ml) and further analysed six days post-infection. To bypass p53 dependent senescence, HDFs were stably infected with a *TP53*-silencing retrovirus followed by subsequent *AR* gene silencing using the protocol described before ^21^.

#### AR overexpression

For the overexpression of *AR*, CAFs were stably infected with a blasticidin-resistant lentivirus that constitutively expresses *AR* (a gift from Karl-Henning Kalland, University of Bergen, Bergen, Norway) or with a lacZ control ^63^. Two days post infection, cells were selected for blasticidin resistance (10 µg/ml). *AR*- or *lacZ*-overexpressing cells were used for further analysis 6 days post-infection.

#### CRISPR/Cas9-mediated gene deletion

All-in-one lentiviral vectors encoding both the RNA guide (gRNA) and the Cas9 protein for *AR* gene deletion were obtained from Applied Biological Materials Inc. (Cat# 12233111). HDFs were infected with high-titer lentiviruses expressing two separate *AR*-targeting gRNAs (#1, #2) or a scrambled control guide RNA. After 48 hours, cells were selected with puromycin (1 µg/ml), and the colonies were pooled before analysis of *AR* disruption. To further confirm the gene targeting efficiency, Surveyor assays were performed one week after the infection. PCR was used to amplify the genomic regions encompassing the two targeted sequences for AR-gRNA. Subsequently, the amplified genomic DNA was then tested for the presence of a DNA-mismatch through incubation with a single-strand-specific endonuclease. The reactions of each targeted sequence were analyzed with a 2% agarose gel and compared to a similar reaction made with control gRNA infected cells for specificity control.

#### Lamin A/C mutant expression

The Q5 Site-Directed Mutagenesis Kit (NEB, Cat# E0554S) was used to generate a Ser301 to Asp (S301D) mutant of lamin A/C from the wildtype lentiviral vector pCDH_blast_MCS_Nard_GFP_LAMIN (Addgene #167340) ^64^. Mutation specific forward and reverse PCR primers, designed using NEBaseChanger tool (https://nebasechanger.neb.com/), were as follows: S301D Forward primer: 5’ CCGCATCGACGACCTCTCTGCCCAG -3’ and S301D Reverse primer: 5’ ATGCGCGACTGCTGCAGC -3’. The resulting mutant LMNA expressing construct (pCDH_blast_MCS_Nard_GFP_LAMIN_S301D) was confirmed by sequencing. The lentiviruses expressing either S301D or wildtype lamin A/C were used to transduce HDFs. Selection with blasticidin (10 µg/ml) was used to create stable cell lines and, six days post infection, the cells were analysed using different assays.

#### Treatments

HDFs and CAFs were treated with different inhibitors or agonists in complete DMEM media containing 10% FBS and 1% antibiotics for a duration of 48-72 hours as specified in figure legends. HDFs were treated with the AR inhibitor AZD3514 (10 µM, Adooq Biosciences, Cat# A12396), the PPP1 inhibitor Tautomycin (10 µM, SantaCruz, Cat# sc-507214) or DMSO as solvent control. CAFs were treated with the AR agonist Ostarine (Selleckchem, Cat# S1174) or DMSO. Furthermore, HDFs were cultured in DMEM supplemented with charcoal-treated FBS for 48 hours prior to treatment with DHT (10 nM, Sigma-Aldrich, Cat# 10300), the AR degrading PROTAC ARCC4 (1 µM, Tocris, Cat# 7254), or ethanol, and treatment was continued for the specified duration.

### Proximity ligation assays

Proximity ligation assays (PLA) were performed using the Duolink® In Situ Red Kit Mouse/Rabbit (Sigma-Aldrich, Cat # DUO92101). For *in vitro* PLA, cells were cultured on 13 mm glass coverslips in 24-well plates for 24 hours and fixed with 4% freshly prepared paraformaldehyde (PFA) for 10 minutes at room temperature (RT). Subsequently, cells were permeabilized with 0.1% Triton X-100 in PBS for 10 minutes at RT, followed by blocking with PLA blocking buffer in a humidified chamber at 37°C for 2 hours. The cells were then incubated overnight at 4°C with the pairs of primary antibodies as indicated in the respective figure legends. The following day, tissue sections were washed three times with the kit-provided buffers, and proximity ligation and rolling cycle probe amplification were performed following the manufacturer’s protocol. Nuclei were counterstained with DAPI (1ug/ml; Merck, Cat# MBD0015).

For tissue PLA, freshly cut frozen tissue sections were fixed in 4% freshly prepared PFA for 10 minutes at RT. Tissue sections were then permeabilized with 0.5% Triton X-100 in PBS for 15 minutes at RT, blocked with PLA blocking buffer in a humidified chamber at 37°C for 2 hours, and incubated overnight at 4°C with pairs of primary antibodies as indicated in figure legends. The following day, tissue sections were washed 3 times with the provided buffers, and proximity ligation and probe amplification were performed following the manufacturer’s protocol. Tissue sections were additionally immunostained with A488-labelled anti-vimentin antibodies (Abcam, Cat# ab185030) and counterstained with DAPI. Details of all the different antibodies and dilutions used in PLA assays are indicated in the Table S1.

Images were acquired using a Zeiss LSM 880 confocal microscope. The number of PLA puncta per cell was quantified using ImageJ’s Analyze Particle tool. Briefly, multi-colour images were split into single channels and converted to grayscale. The threshold was adjusted to highlight structures of interest, and background pixels were subtracted. The ’Watershed’ option was used to separate merged nuclei. The binary image was then analysed using the ’Analyze Particles’ function, with the settings adjusted to outline the nuclei. The same steps were repeated to analyze the image corresponding to PLA complexes. The outline images corresponding to nuclei and PLA complexes were inverted, resulting in an image where selected nuclei appeared in blue and PLA complexes in red. The PLA complexes outside the selected nuclei were excluded, and the average number of PLA puncta per cell was calculated.

### Immunofluorescence

Frozen tissue sections or cultured cells on glass coverslips were fixed with cold 4% PFA for 15 minutes at RT. The fixed cells were washed with PBS and permeabilized with 0.1% Triton X-100 in PBS for 10 minutes, followed by incubation with 2% bovine serum albumin (BSA) in PBS for 2 hours at RT. Primary antibodies were diluted in 2% BSA in PBS (pH 7.6) and incubated overnight at 4°C. After washing three times in PBS, the samples were incubated with fluorescently labeled secondary antibodies (Invitrogen) for 1 hour at RT. Following washing with PBS, slides were mounted with Fluoromount Mounting Medium (Sigma-Aldrich) after DAPI staining. Details of antibodies and dilutions used for immunostaining are indicated in the Table S1.

Immunofluorescence images were acquired using a ZEISS LSM880 confocal microscope with 20X, 40X or 63X objectives, and ZEN black software was used for image acquisition and processing. For fluorescence signal quantification, acquired images for each color channel were analyzed using ImageJ software, using the “measurement” or “particle analysis” functions to select areas or cells of interest (tumor or stromal). All measurements were exported as Microsoft Excel data files and data were plotted using GraphPad Prism 9.5.1 software. Imaris software v9 (Oxford Instruments) was used to create the 3D images. Z stack images acquired with Zeiss LSM880 confocal laser scanning microscope was utilized to recreate the volume at full resolution. The surface rendering method was used to create the 3D models. The surface area detail level (grain size) was set at 0.24 µm, with an upper background subtraction of 0.904 µm and a lower background subtraction threshold value of 10. The images were exported as TIFF.

### Transmission electron microscopy (TEM)

The sample preparation and TEM analysis were carried out at the Electron Microscopy Facility at the University of Lausanne. Briefly, cells were grown on glass coverslips and fixed by incubating them with a fresh mixture of 2.5% glutaraldehyde (Sigma-Aldrich, Cat# G5882), 1% osmium tetroxide (EMS, Cat# 19110), and 1.5% potassium ferrocyanide (Sigma-Aldrich, Cat# 60279) in 0.1 M phosphate buffer (pH 7.4) for 1 hour at room temperature. The samples were washed three times with distilled water, and dehydration was performed by immersing the coverslips in 30% acetone for 5 minutes, followed by 50% acetone for 5 minutes. Subsequently, the coverslips were treated with a 0.3% uranyl acetate solution in 50% acetone for 20 minutes. The coverslip was then rinsed with a 70% acetone solution in water for 5 minutes, and three sequential 5-minute washes were performed using 100% acetone. A drop of 100% Epon resin was added to the coverslips and left overnight in a fume hood. The next morning, a gelatin capsule was completely filled with 100% Epon resin, turned over, and placed on the coverslip. The specimen was then polymerized at 60°C for 48 hours. Ultrathin sections of 50 nm were cut on a Leica Ultracut (Leica) and transferred onto a copper slot grid (2 × 1 mm; EMS) coated with a polystyrene film. The sections were post-stained with 2% uranyl acetate (Sigma-Aldrich) in H2O for 10 minutes, followed by rinsing with H2O. Finally, the sections were stained with Reynolds’s lead citrate (Sigma-Aldrich, Cat# 15326) in H2O for 10 minutes and rinsed for a final time with H2O. Micrographs were taken with a transmission electron microscope Philips CM100 (Thermo Fisher Scientific) at an acceleration voltage of 80kV, using a TVIPS TemCam-F416 digital camera (TVIPS).

### Cancer cells expansion assays

Cancer cell expansion assays were conducted as previously described ^21^ with minor modifications. SCC13 tumour cells and fibroblasts were mixed in a 1:1 ratio in complete DMEM containing 1% Matrigel (BD Biosciences, Cat# 354230). Cells were plated on 8-well glass bottom culture slides (Corning, Cat# 354118), precoated with a thin layer of 10% Matrigel. Cells were allowed to grow for four days under normal culture conditions before fixing with 4% PFA. Immunofluorescence staining of HDFs and SCC13 cells were carried out with anti-vimentin and anti-pan-keratin antibodies and counterstained with fluorescently labelled secondary antibodies, as described above. Each well on the slides was imaged using a Zeiss LSM880 confocal microscope with a 20X objective, and the percentage of SCC-positive area was quantified from five random fields per well using ImageJ software. The expansion of tumour cells was represented as a percentage of the SCC13 positive area.

### 3D Spheroid Assay

8-well glass bottom culture slides (Corning, Cat# 354118) were coated with 80% Matrigel (100 μl per well; BD Biosciences, Cat# 354230) and incubated at 37°C for 30 minutes to allow polymerization. RFP-expressing SCC13 cells and fibroblasts were mixed 1:1 in complete DMEM supplemented with 1% Matrigel and overlaid on wells containing polymerized Matrigel. The spheroids were allowed to grow under normal culture conditions for six days. For immunostaining, the spheroids were fixed with 4% PFA and 2.5% glutaraldehyde for 10 minutes, carefully washed with PBS and stained with fluorescently labeled anti-vimentin antibody (Abcam, Cat# ab185030). Images of the spheroids were captured using a Zeiss LSM880 microscope with a 10X objective, and the diameter and number of spheroids were determined using ImageJ analysis.

### Western blotting

Cells were lysed for 30 minutes on ice in lysis buffer (150 mM NaCl, 1.0% NP-40, 0.5% sodium deoxycholate, 0.1% SDS, 50 mM Tris, pH 8.0) containing a 1X protease/phosphatase inhibitor cocktail (Thermo Scientific, Cat# 78440). Protein concentration was then determined using a Pierce BCA protein detection kit (Thermo Scientific) and samples were normalised to 2 µg/1ul. Proteins were then mixed with SDS-PAGE loading buffer (Tris pH 7.5 20mM, EDTA 1mM, SDS 1%) and boiled at 95°C for 10 minutes. Proteins were then separated on 10-12% SDS-PAGE gels and transferred to a polyvinylidene difluoride (PVDF) membrane using the Trans Blot Turbo™ transfer system (Bio-Rad). Membranes were blocked and primary antibodies were diluted in 2% BSA in Tris-buffered saline (TBS). The detection was performed by the use of peroxidase-conjugated secondary antibodies with the SuperSignal West Pico (Thermo Scientific). Signals were finally detected on Fuji Medical X-ray films (Fujifilm) or using iBright imaging systems (Invitrogen).

### Time-lapse imaging and NERDI dynamics

GFP protein fused with a nuclear localization signal (GFP-NLS) was stably transduced into HDFs using lentiviruses (pTRIP-SFFV-EGFP-NLS; Addgene, Cat# 86677). Cells were cultured on eight-well coverslip-bottomed slides (ibidi GmbH, Cat# 80827), and imaged using a Zeiss Observer Z1 microscope equipped with a temperature- and CO2-controlled stage-top incubation unit (Tokai Hit). Images were acquired at the indicated time intervals using an AxioCam MRM CCD camera (Zeiss) with a 20X objective. To minimize photodamage and phototoxicity, the illumination was set to 10% with an exposure time of 80 ms per slice.

For the analysis of nuclear envelope rupture during interphase (NERDI), time-lapse imaging was performed with 10-minute intervals. The mean fluorescence intensity of GFP– NLS (BacMam, Thermo Fisher) within a defined region of the cell nucleus was quantified using ImageJ software. For data normalization, the peak fluorescence intensity of each sample was set to 1, and the lowest fluorescence intensity was set to 0. The data were aligned around the time point of NERDI and plotted using GraphPad Prism 9.5.1.

### Co-Immunoprecipitation

HDF cells were transduced with lentiviruses expressing either control or *AR*-targeting shRNA and selected with puromycin. After six days, cells were harvested and counted. Nuclear extracts were prepared from an equal number of control- or *AR* silenced cells using the NE-PER™ kit (Thermo Scientific, Cat# 78833), according to the manufacturer’s protocol. Briefly, the cell pellet was resuspended in ice-cold CER-I buffer containing a 1X protease/phosphatase inhibitor cocktail (Thermo Scientific, Cat# 78440). After vortexing for 15 seconds and incubating on ice for 10 minutes, the required volume of ice-cold CER II buffer was added. The pellet was vortexed for a further five seconds and centrifuged at maximum speed for five minutes. The supernatant containing the cytoplasmic extract was discarded and the pellet containing the nuclei was then lysed in ice-cold NER buffer containing a 1X protease/phosphatase inhibitor cocktail, vortexed for 15 seconds, and incubated on ice for 40 minutes with intermittent vortexing. The nuclear lysates were sonicated briefly with five pulses, and centrifuged at maximum speed for 10 minutes. The pellet containing chromatin and insoluble nuclear components was discarded, while the supernatant containing the nuclear extract was stored on ice, quantified and used for Co-IP. Equal amounts of nucelar lysates from control- and *AR* silenced cells were then diluted with Co-IP buffer (20 mM Tris-HCl pH=8, 200 mM NaCl, 0.2% NP4O, 2% glycerol, 0.05% sodium deoxycholate, 1.5 mM MgCl2, 1X protease/phosphatase inhibitor cocktail) to a volume of 1 ml. The samples were then precleared with 10 ul of Protein A Dynabeads (Invitrogen) and 1 ul of DNase for 30 minutes at RT. Beads were removed and nuclear extracts were incubated overnight at 4°C with either anti-lamin A/C antibody (Abcam, #ab108595) or non-immune IgG. The following day, preequilibrated Protein A Dynabeads were added, and the samples were incubated for at least two hours at RT on a rotator. The supernatant was removed using magnetic separation, and the beads bound with the protein of interest were washed three times with 1 ml of 0.5X Co-IP buffer for 10 minutes at RT. Finally, the beads were washed with 1X PBS and subjected to mass spectroscopy or immunoblot experiments.

### Proteomics analysis

The proteomics analysis was performed at the Protein Analysis Facility of the University of Lausanne using the following methods.

#### SDS-PAGE and in-gel digestion

Co-IP samples were loaded onto a 12 % mini polyacrylamide gel and migrated approximately 2-3 cm before being stained by Coomassie blue. Gel lanes between 10-300 kDa were excised into five-six pieces and subjected to in-gel digestion using sequencing-grade trypsin (Thermo Scientific, Cat# 90057), as previously described ^65^. The extracted tryptic peptides were then dried and resuspended in a 0.05% trifluoroacetic acid and 2% (v/v) acetonitrile solution.

#### Liquid Chromatography-Mass Spectrometry analyses

Tryptic peptide mixtures were injected into a Dionex RSLC 3000 nanoHPLC system (Dionex, Sunnyvale, CA, USA), coupled with a high resolution QExactive Plus mass spectrometer (Thermo Fisher) via a nanospray Flex source. Peptides were loaded onto a trapping microcolumn (Acclaim PepMap100 C18; 20 mm x 100 μm ID, 5 μm, Dionex) and separated on a custom-packed C18 column (75 μm ID × 45 cm, 1.8 μm particles). A gradient ranging from 4 to 76 % acetonitrile in 0.1 % formic acid for peptide separation (total time: 65 minutes). Full MS survey scans were performed at a resolution of 70,000. In data-dependent acquisition controlled by Xcalibur software (Thermo Fisher), the 10 most intense multiply charged precursor ions detected in the full MS survey scan were selected for higher energy collision-induced dissociation (HCD, normalized collision energy NCE=27 %) and analysis in the orbitrap at a 17’500 resolution. The window for precursor isolation was of 1.5 m/z units around the precursor and selected fragments were excluded for 60 s from further analysis.

#### Data processing

MS data were analyzed using Mascot 2.7.0 (Matrix Science, London, UK) with a search against the SwissProt database, restricted to the human (Homo sapiens) taxonomy (www.uniprot.org, version of June 2020, containing 20’746 sequences), and a custom contaminant database containing common environmental contaminants and digestion enzymes. Trypsin (cleavage at K,R) was used as the enzyme definition, allowing for up to 2 missed cleavages. Mascot searches were performed with a parent ion tolerance of 10 ppm and a fragment ion mass tolerance of 0.02 Da. Carbamidomethylation of cysteine was specified as a fixed modification, while protein N-terminal acetylation and methionine oxidation were specified as variable modifications.

Scaffold software (version Scaffold 4.11.1 or 5.0.0, Proteome Software Inc., Portland, OR) was used to validate MS/MS-based peptide and protein identifications by Mascot. Peptide identifications were accepted if they achieved a probability greater than 90.0% using either the Scaffold Local FDR algorithm (v. 4.11.1) or the Percolator posterior error probability calculation (v. 5.0.0) ^66^. Protein identifications were accepted if they had a probability greater than 95.0% and contained at least two identified peptides. Protein probabilities were determined using the Protein Prophet algorithm ^67^. Proteins that contained similar peptides and could not be differentiated based on MS/MS analysis alone were grouped to satisfy the principles of parsimony. Proteins sharing significant peptide evidence were grouped into clusters.

For the analysis of LMNA phosphorylation, the MS data from the corresponding gel bands were subjected to a secondary Mascot search, incorporating phosphorylation of serine, threonine and tyrosine as additional variable modifications. MsViz software ^68^ was used to compare intensities of phosphorylation sites between samples.

### Co-IP with recombinant proteins

I*n vitro* protein interactions with recombinant proteins were performed as follows. 200 ng purified Myc-tagged lamin A (MW: 74 KDa, OriGene; Cat# TP304970) and 200 ng 6XHis tagged AR (1-556 aa, 63 kDa, Ray Biotech; Cat# RB-14-0003P) were mixed in 500 µl binding buffer (150 mM NaCl, 20mM Tris-HCl; pH-8, 0.1% NP40, 10% glycerol) and incubated overnight at 4°C on a rotary platform. The next day, 20 µl of protein A magnetic beads pre adsorbed with 0.5 µg of anti-lamin A/C antibody (Abcam, #ab108595) were added to the protein mixture and incubated for 2 hours at RT. Bead-bound protein complexes were separated from the unbound fraction by magnetic pulldown, beads were washed 3 times in binding buffer and then resuspended in 30 µl of SDS-PAGE loading buffer. As a control for specificity, parallel pull-down reactions were performed with AR protein previously denatured by a short heat treatment (20 min at 85°C), or the protein mixture was incubated in high salt buffer (800 mM NaCl). Immunoprecipitates with anti-lamin A/C antibody were analysed in parallel with 5% unbound supernatants by immunoblotting with anti-lamin A/C (Cell Signaling; Cat# #4777) or anti-His (Cell Signaling; Cat#12698) antibodies.

### ChIP-seq

Control- and AR-silenced fibroblasts were cross-linked with 1% formaldehyde for 10 minutes at RT, followed by quenching with glycine (final concentration 125 mM). After washing with ice-cold PBS, cells were collected through centrifugation (400g). For the analysis of total lamin A/C chromatin binding, cells were lysed and chromatin was prepared using the Chromatin EasyShear Kit - High SDS (Diagenode; Cat# C01020012), and shearing of the chromatin was achieved using the Bioruptor with four rounds of ten cycles of 30 seconds “ON„ and 30 seconds “OFF„ at HIGH power setting (position H). To determine chromatin binding of pSer301 and wild-type lamin A/C together with histone markers, chromatin was prepared using the Chromatin EasyShear Kit - Low SDS (Diagenode, Cat# C01020013) and sonicated using an E220 focused ultrasonicator (Covaris).

The ChIP reaction was performed using the iDeal ChIP-seq kit (Diagenode, Cat# C01010055). Samples were pre-cleared using the beads provided in the kit and incubated overnight at 4°C with 5 µg of commercially available antibodies against lamin A/C (Abcam, #ab108595), phospho-Ser301 lamin A/C (Affinity, #AF7177), H3K27Ac (Abcam, #ab4729), H3K9Me3 (Abcam, #ab8898). Antibody-chromatin complexes were pulled down using protein A-coated magnetic beads. Elution was performed according to the manufacturer’s instruction and chromatin was quantified using the Qubit Fluorometric Quantification Kit (Thermo Scientific). For library preparation, 10 ng of DNA was utilized with the NEBNext® ChIP-Seq Library Prep Reagent Set for Illumina. Subsequently, pair-end sequencing was performed at the Novegene facility using the Novaseq 6000 platform.

The FASTA files were aligned using the Burrows-Wheeler Aligner (BWA) ^69^ and narrow peaks were detected using MACS2 software ^70^ with a q-value cut-off of 0.05. Graphic illustrations of ChIP-seq peaks were generated using the Easeq tool, an interactive software for analysis and visualisation of ChIP-sequencing data (https://easeq.net/). Peaks were annotated and merged using the annotatePeaks.pl and mergepeaks.pl functions available in the HOMER software ^71^.

To detect lamin A/C associated with large genomic regions (LADs), we used the Enrichment Domain Detector (EDD), a tool developed to analyse broad enrichment domains from ChIP-seq data ^42^. Mapped reads from BAM files were used to call LADs, with automatic estimation of GapPenalty. Log ratios between input and ChIP samples were calculated using a BinSize of 10 Kb, with the human genome hg38 as the reference assembly. Peaks were retrieved with an FDR p-value of 0.05.

### ATAC-seq

ATAC-seq analysis was conducted on the same batch of *AR*-silenced and control HDFs that were used for ChIP-seq analysis. Viable cells were harvested, and their viability was assessed using the trypan blue assay and cell counting. ATAC-seq was performed as previously reported ^41, 72, 73^. Briefly, nuclei were extracted from 50,000 cells and the nuclear pellet was resuspended in the Tn5 transposase reaction mix (Diagenode, Cat# C01070012-30). The transposition reaction was incubated at 37°C for 30 min. Afterwards, DNA fragments were purified and isolated using the MinElute Reaction Cleanup Kit (Qiagen, Cat # 28204). The purified DNA fragments were quantified using Qubit, and library preparation was performed. ATAC-seq library preparation involved PCR amplification of the purified DNA fragments using Diagenode primer indices (Diagenode, Cat# C01011034). Amplification cycles were adjusted and libraries were purified using AMPure beads. Library quality was assessed using Qubit and Bioanalyzer, and the final libraries were sequenced on an Illumina Novaseq 6000 machine.

For ATAC-seq data analysis, raw data underwent quality control using FastQC. Trimming and filtering were performed, which included the removal of adapter sequences, low-quality reads (average quality score below 20, an N ratio exceeding 15%, adapters, or a length of less than 18 nucleotides after trimming), and trimming of adapter sequences at the 3’ end. The reads were then aligned to the human reference genome (GRCh38/hg38) using BWA ^69^. Following alignment, peaks were called using the MACS2 algorithm (http://liulab.dfci.harvard.edu/MACS). To analyze the distribution of peaks across different functional domains, ChIPseeker software ^74^ was used.

### Transcriptomic analysis

RNA extraction and purification was carried out using the Direct-zol RNA Miniprep Kit (Zymo Research). The quality of RNA samples was assessed, ensuring a purity of OD260/OD280 ≥ 1.8 and RIN ≥ 8. Global transcriptomic analysis was conducted at the Institute of Genetics and Genomics of Geneva (iGE3) using Clariom™ D GeneChip assays. Briefly, the GeneChip® WT PLUS Reagent Kit (Thermo Fisher Scientific, Cat. No. 902280) was used to prepare samples and to facilitate hybridization to the human Clariom™ D arrays (Thermo Fisher Scientific). Transcriptome Analysis Console (TAC) software (Thermo Fisher Scientific) was used for data processing and analysis.

Gene set enrichment analysis (GSEA) for Clariom D array expression profiles was performed as in ^21^ by using GSAA-SeqSP software (gene set association analysis for array expression data with sample permutation) from the Gene Set Association Analysis (GSAA) platform (version GSAA_2.0, http://gsaa.unc.edu/). Curated gene sets were obtained from the Molecular Signatures Database (MSigDB v5.2; http://www.broadinstitute.org/gsea/msigdb/) or from the GEO database with accession numbers as indicated.

Normalised single-cell RNA-seq profiles generated from head and neck squamous cell carcinoma (HNSCC) samples using the 10X Genomics platform were downloaded from Gene Expression Omnibus (GSE188737) ^50^. Scaling and dimensionality reduction with PCA and UMAP were performed using Seurat v4.3.0 (www.satijalab.org/seurat). The top 200 up regulated genes from the Lamin A/C S301D transcriptomic data were used to calculate signature scores using the Seurat function ’AddModuleScore’ 75. All analyses were performed in R v4.2.1.

RT-qPCR analysis was performed as described previously ^21^. A list of primers used in this study is provided in the Table S1.

### Tumorigenesis experiments

Mouse back injections were carried out in 9-week-old NOD/SCID male mice (with IL-2 receptor γ chain null mutation, *NOD.Cg-Prkdcscid Il2rγtm1Wjl/SzJ*, (the Jackson Laboratory). 2.5 x 10^5^ RFP-SCC13 tumor cells, admixed with an equal number of HDFs expressing either S301D mutant or wildtype lamin A/C were re-suspended in 70 µl of Matrigel solution (BD Bioscience) and injected intradermally into the left and right side of the mouse back. Mice were sacrificed for tissue analysis 14 days after injection. The tumor volumes were calculated using the formula V = 3.14 × (W2 × L)/6 V= volume, W= width, and L= length. Animal experiments were completed in accordance to the Swiss guidelines and regulations for the care and use of laboratory animals with approved protocol from the Canton de Vaud veterinary office (animal license No. 1854.4e).

### Statistical analysis

Statistical testing is performed using GraphPad Prism 9.5.1 Software. Data are presented as mean ± SE, as indicated in the legend. For each experiment, two to three separated cell strains were used in independent experiments. Statistical significance for comparing two experimental conditions is calculated by two-tailed unpaired t-tests, while for comparing one control with two experimental conditions the statistical significance is calculated using the non-parametric one-way ANOVA test. All P values < 0.05 are considered statistically significant.

