## Supplemental Figure S1-S7 and Table S1 for "Nuclear lamin A/C phosphorylation by loss of Androgen Receptor is a global determinant of cancer-associated fibroblast activation"

**Figure S1** (related to Fig. 1)

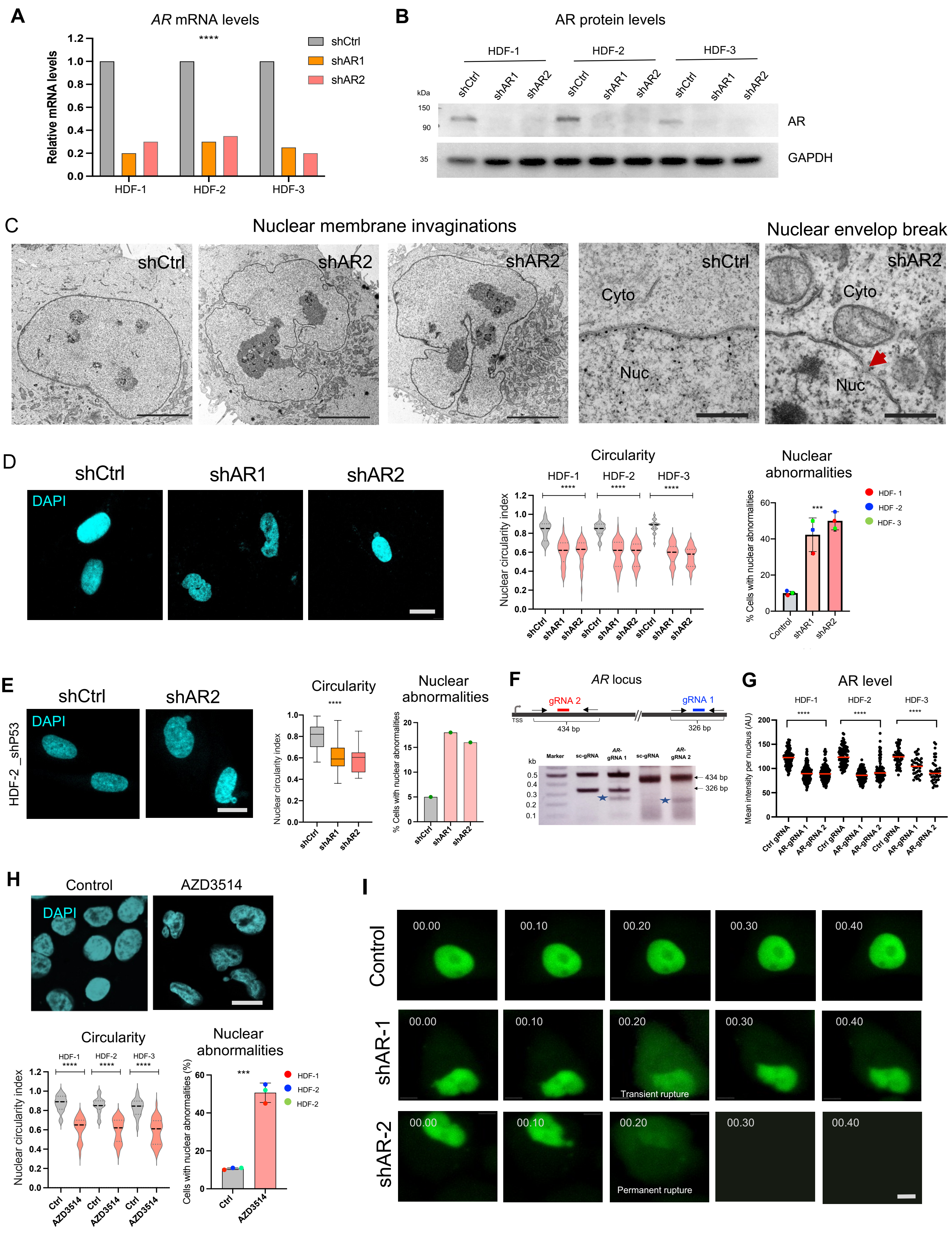

**Figure S1**. **Nuclear membrane abnormalities resulting from loss of AR in HDFs.**

A) RT-qPCR analysis of *AR* mRNA levels in three HDF strains infected with two *AR* silencing lentiviruses (shAR1, shAR2) versus control (shCtrl). The statistical significance was determined using values of the three strains plus/minus *AR* silencing together, by non-parametric one-way ANOVA test, n=3 strains; ****P<0.0001.

B) Immunoblot analysis of AR protein levels in the same HDF strains plus/minus *AR* silencing as in previous panel, with GAPDH as equal loading control.

C) Transmission electron microscopy images of nuclei from control and *AR*-silenced HDFs. Shown are low and high magnification images with nuclear membrane (NM) invaginations and nuclear envelope breaks (arrowheads) in *AR* silenced cells. Scale bar : 5 μm, left three panels; 0.4 μm, right two panels.

D) Nuclear alterations of HDFs cultured on collagen-coated soft matrix upon *AR* silencing. Shown are representative images of DAPI-stained nuclei and quantification of nuclear circularity index and nuclear abnormalities in three HDF strains infected with two *AR* silencing lentiviruses (shAR1, and shAR2) versus control (shCtrl). Cells were cultured on collagen-coated cover glasses for 48 h. Measurements were taken as in Fig. 1E,F. For circularity, n=>100 cells per condition; for abnormalities, n=3 HDF strains; ****P<0.0001, ***P<0.001, non-parametric one-way ANOVA Kruskal–Wallis test, mean±SE. Scale bars: 10 μm.

E) Nuclear alterations resulting from *AR* silencing in HDFs with silenced *TP53*. Shown are representative images and quantification of nuclear circularity index and nuclear abnormalities in HDFs stably infected with an *TP53* silencing retrovirus and subsequently infected with two *AR* silencing versus control lentiviruses. For circularity, n=>50 cells per condition; ****P<0.0001, non-parametric one-way ANOVA Kruskal–Wallis test, mean±SE.

F) Surveyor Assay of HDFs, one week after the infection with lentiviruses co-expressing the Cas9 enzyme and one of two RNA guides targeting *AR* (*AR*-gRNA1 and *AR*-gRNA2) or a scrambled control gRNA (sc-gRNA). The two targeted genomic regions of the *AR* gene (#1,2) were tested for DNA mismatch by PCR with corresponding oligo pairs at the indicated position in the presence of single-strand specific endonuclease. Shown are the amplified PCR products of the expected size (arrows) and endonuclease cleavage products (asterisks).

G) Quantification of AR florescence signal intensity in three HDF strains infected with the two *AR*-targeting versus control CRISPR lentiviruses as in the previous panel. Representative images of IF analysis with anti-AR antibodies are shown in Fig. 1E. Quantification of IF fluorescence signal intensity per cell was carried out as described for Fig. 2C. n=>50 cells per condition, ****p<0.0001, non-parametric one-way ANOVA Kruskal–Wallis test.

H) Nuclear alterations in three HDF strains plus/minus treatment with the AR inhibitor AZD3514. HDFs were treated with either AZD3514 (10 μm) or DMSO control (Ctrl) for 48h followed by DAPI staining. Shown are representative images and quantification of nuclear alterations as in Fig. 1F. For circularity, n=>50 cells per condition; for abnormalities, n=3 HDF strain (>50 cells per condition); ****P<0.0001, ***P<0.001, non-parametric one-way ANOVA Kruskal–Wallis test, mean±SE. Scale bar : 10 μm.

I) Representative snapshots of time-lapse imaging analysis of HDFs plus/minus *AR* gene silencing and expression of a green fluorescence protein with nuclear localization signal (GFP–NLS) as described in Fig.1G. Images shown here depict transient nuclear envelope rupture (second row; middle image) and permanent nuclear rupture (third row; middle image) events in HDFs with silenced *AR*. Time is shown on the top of each panel (hours: minutes). Scale bar: 10 μm.

### **Figure S2** (related to Fig.2)

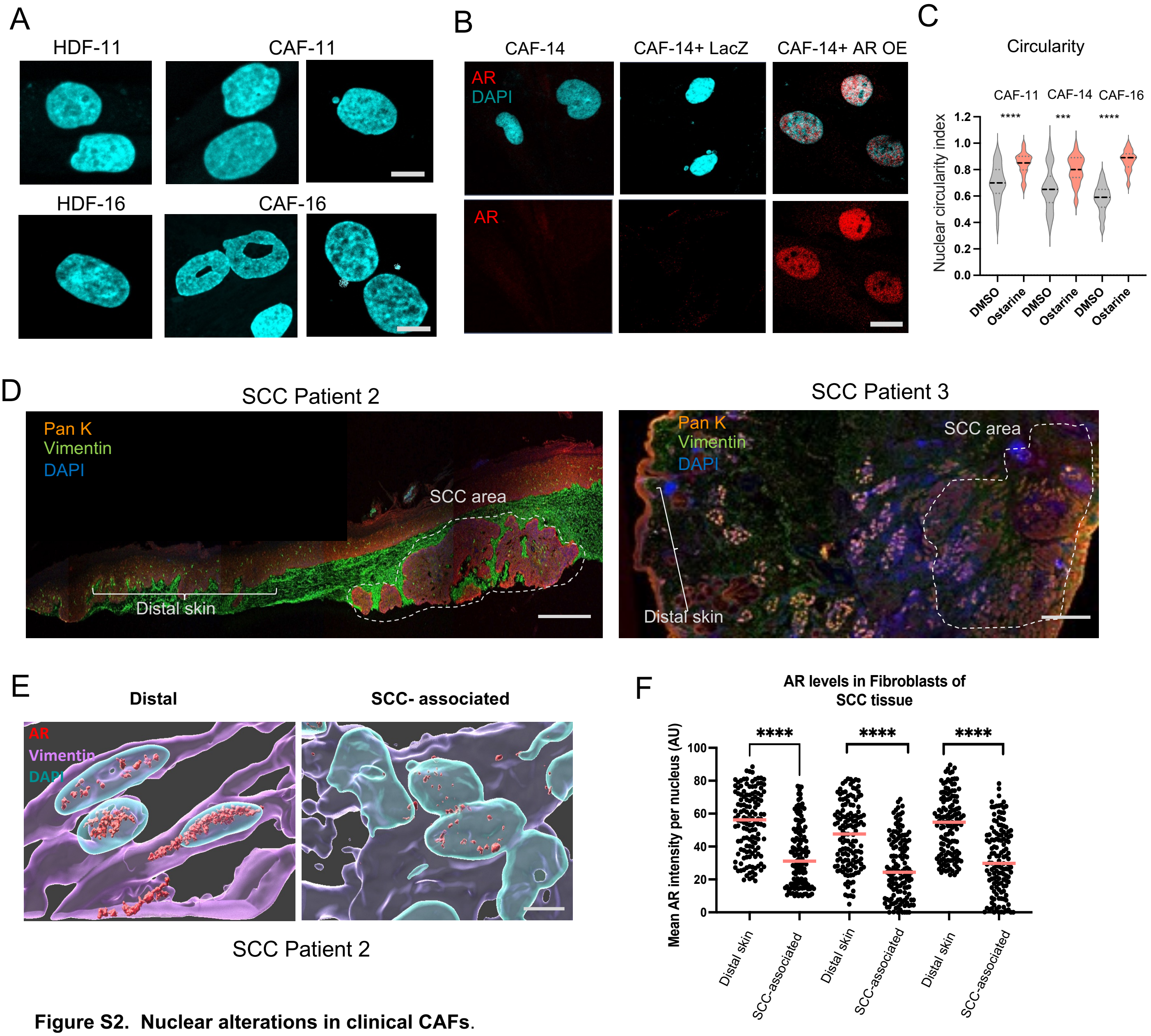


**Figure S2. Nuclear alterations in clinical CAFs.**

A) Representative images of DAPI-stained nuclei of an additional pair of CAFs and matched HDFs as shown and quantified in Fig. 2A. Scale bar: 5 µm.

B) Representative images of a CAF strain infected with a LacZ control (LacZ) or *AR* overexpressing lentivirus as in Fig. 2B, analyzed by IF with anti-AR antibodies and DAPI staining. Quantifications are provided in Fig. 2B. Scale bar: 10 µm.

C) Quantification of circularity index of three pairs of CAFs treated with the AR agonist Ostarine (10 µM) or DMSO control for 48 hours. Quantification was carried out as in Figs. 1B and 2A. n=>100 cells per condition; ***P<0.001, **P<0.01; two-tailed unpaired t-test.

D) Immunofluorescence analysis of two additional skin SCC lesions with anti-pan-keratin (Red) and -Vimentin (red) antibodies besides the lesion shown in Fig. 2E. The SCC and flanking unaffected areas utilized for further analysis are marked with dotted and solid lines, respectively. Scale bar 100 µm.

E) Representative 3D surface reconstruction images from another SCC and flanking skin besides those shown in Fig. 2G of DAPI-stained nuclei (cyan) of vimentin-positive fibroblasts (violet) concomitantly stained with anti-AR antibodies (red). Scale bar: 5 µm.

F) Quantification of anti-AR fluorescence signal intensity in vimentin-positive cells (fibroblasts) in SCC lesions (SCC-associated) versus distal unaffected skin (distal) quantified for nuclear abnormalities in Fig. 2H,I. Quantification of AR fluorescence signal intensity assessed by Image J and expressed in arbitrary units (AU). n(cells)>50 per condition, ****p<0.0001, paired t-test with Mann-Whitney correction.

### Figure S3 (related to Fig.3)

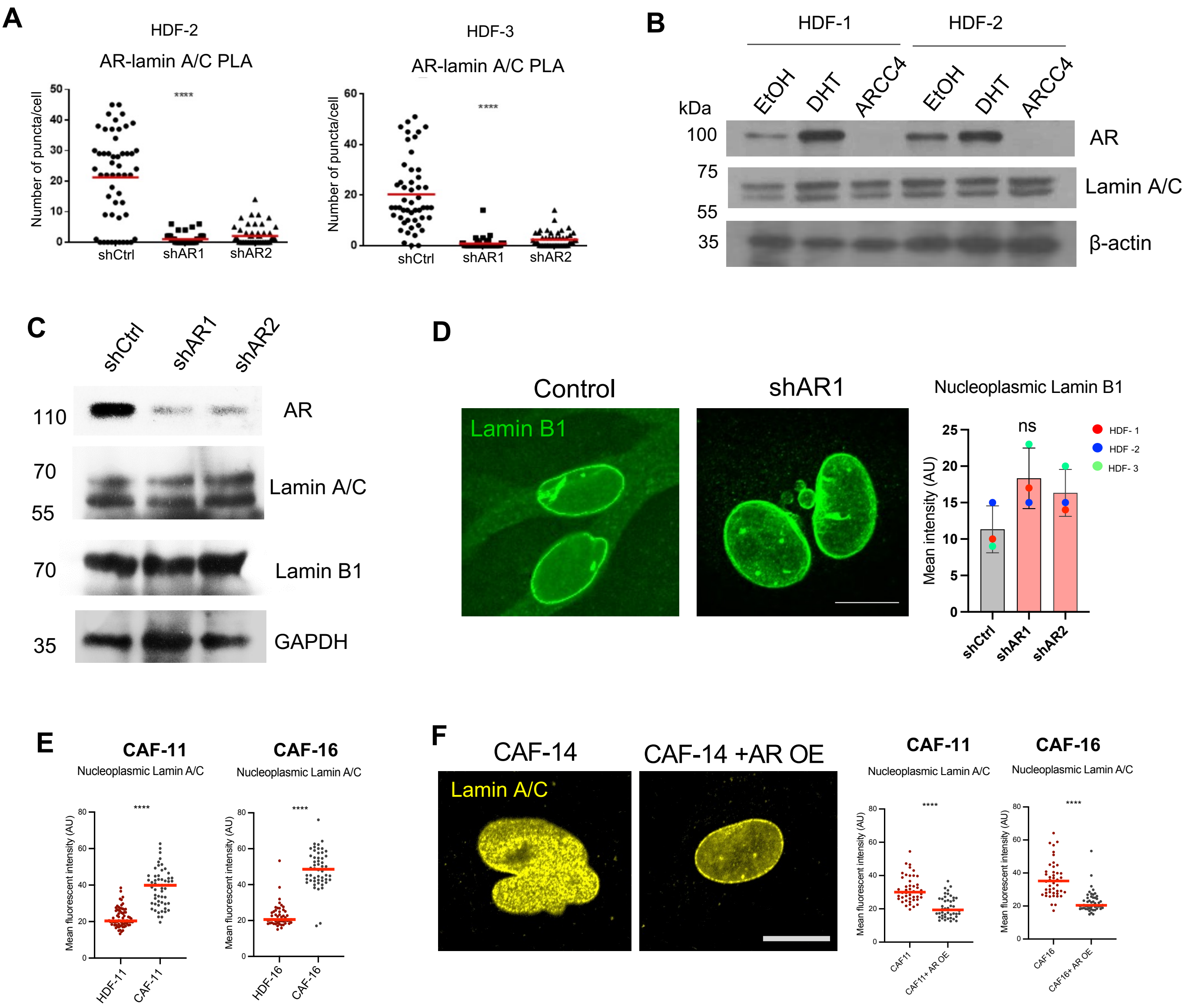


**Figure S3. AR is a determinant of nuclear lamin A/C localization.**

A) Proximity ligation assays (PLA) with antibodies against AR and lamin A/C of two HDF strains, with *AR* silencing as specificity control. Shown is the quantification of the number of puncta per cell together with mean. n(cells)>50 per condition; ****p<0.0001; non-parametric one-way ANOVA test.

B) Immunoblot analysis with anti-AR and lamin A/C antibodies and β-actin, as equal loading control, of two HDF strains treated with DHT (10 nM), ARCC4 (1 µM) or ethanol vehicle control for 72 hours. Cells were cultured in medium supplemented with charcoal-stripped FBS.

C) Immunoblot analysis of AR, lamin A/C, and lamin B1 protein levels in HDFs infected with two *AR* silencing lentiviruses versus control. GAPDH was probed as a loading control.

D) Lamin B1 expression and localization in HDFs plus/minus *AR* silencing. Shown are representative IF images and quantification of fluorescence signal intensity of nucleoplasmic lamin B1, determined by Image J and expressed in arbitrary units (AU) per nucleus. The data shown are the mean ± SE. >100 cells per condition. n=3 HDF strains. No significant difference was observed with the non-parametric one-way ANOVA test. Scale bar 5 µm.

E) Quantification of nucleoplasmic lamin A/C in CAFs and matched HDFs derived from the same patients. The lamin A/C fluorescence signal intensity in arbitrary units (AU) per nucleus (dots) is shown together with mean and statistical significance. n=>50 cells per condition; ****P<0.0001; two-tailed unpaired t-test.

F) Representative images and quantification of nucleoplasmic lamin A/C in two CAF strains plus/minus AR overexpression. The lamin A/C fluorescence signal intensity in arbitrary units (AU) per nucleus (dots) is shown together with mean. n=>50 cells per condition. ****P<0.0001; two-tailed unpaired t-test. Scale bar 10 µm.

**Figure S4** (related to Fig. 4)

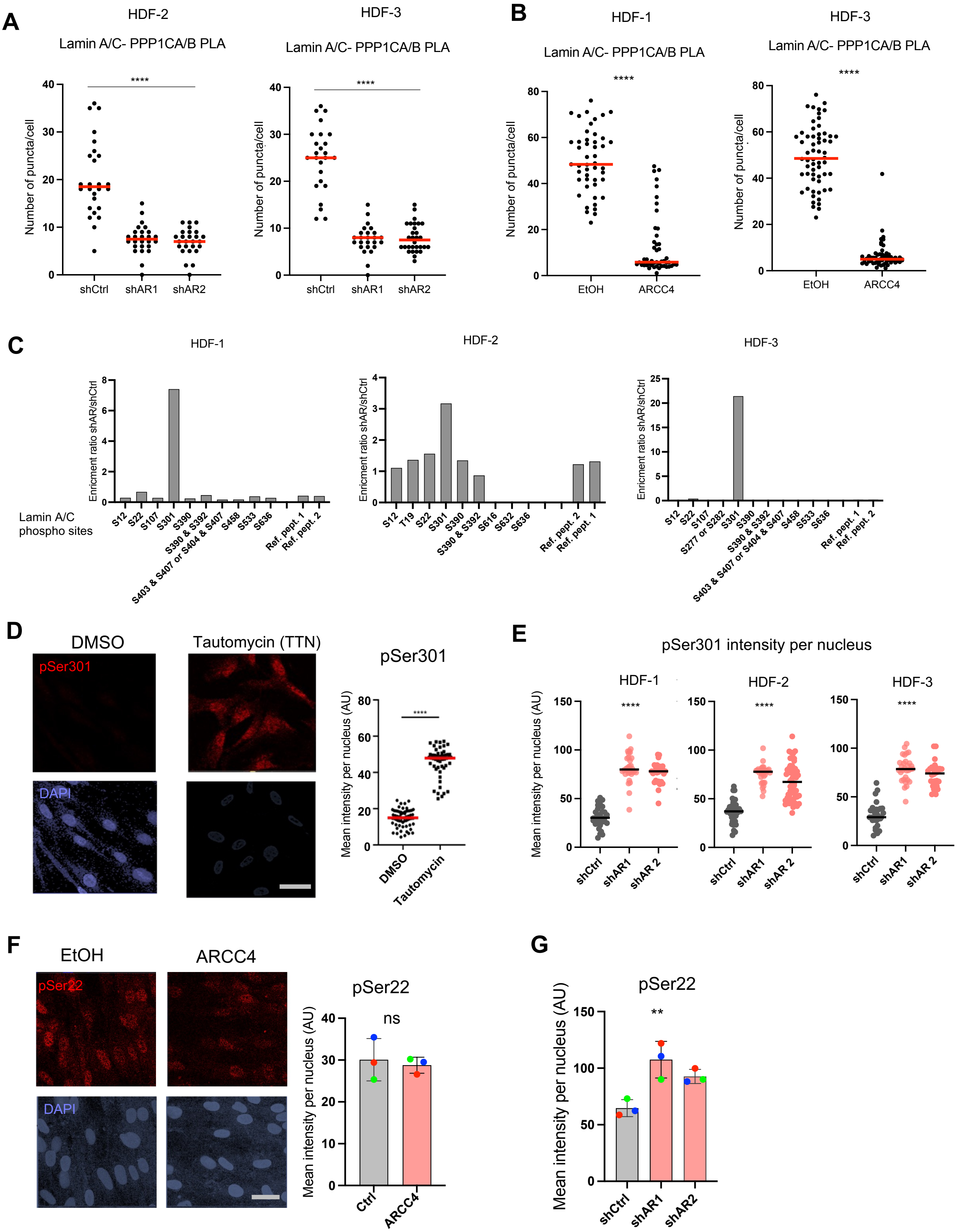

**Figure S4. Loss of AR results in decreased lamin A/C – PPP1 association and increased lamin A/C phosphorylation at Ser 301.**

A) Quantification of PLA signal for lamin A/C and PPP1CA/B association in two different HDF strains infected with *AR* silencing or control viruses. n(cells)>30 per condition, ****p<0.0001, non-parametric one-way ANOVA Kruskal–Wallis test.

B) Quantification of PLA signal for lamin A/C - PPP1CA/B association in two additional HDF strains besides that shown in Fig. 4D treated with the AR degrading compound ARCC4 (1 μM) or ethanol control for 72 h. n(cells)>30 per condition, ****p<0.0001, unpaired student t-test.

C) Enrichment of phospho sites in MS anaylsis of lamin A/C immunoprecipates from three HDF strains with AR silencing vs control. A relevant peptide was selected for each phosphosite and quantified based on signal intensity (precursor mass intensity). Two reference peptides were also measured to evaluate difference in lamin A/C amounts in samples. The enrichment ratio was calculated based on the intensity of a specific peptide in shAR over control.

D) Immunofluorescence analysis of HDFs treated with the PPP1 inhibitor tautomycin (TTN;10 μM) or DMSO vehicle alone for 48 h with antibodies against phospho-Ser301-lamin A/C (pSer301; red) and DAPI for nuclear staining. Shown are representative images and quantification of the anti-pSer301-lamin A/C signal using image J. n(cells)>50 per condition, ****p<0.0001; unpaired student t-test. Scale bar: 20 μm.

E) Quantification of the anti-phospho-Ser301 lamin A/C IF signal intensity per individual cells (dots) in three different HDF strains plus/minus AR silencing. Representative images and overall quantification per strain are shown in Fig. 4G. n(cells)>50 per condition, ****p<0.0001; non-parametric one-way ANOVA test.

F) Immunofluorescence analysis of three HDF strains, treated with ARCC4 (1 μM) or ethanol vehicle alone (EtOH) for 72 h, with antibodies against phospho-Ser22 lamin A/C (pSer22; red) with DAPI for nuclear staining. Shown are representative images and quantification of phospho-Ser22 lamin A/C fluorescence signal intensity assessed by Image J and expressed in arbitrary units (AU). n= 3 strains (>50 cells per condition), no significant difference was observed with the unpaired student t-test, mean ± SE. Scale bar: 20 μm.

G) Quantification of immunofluorescence signal intensity of three HDF strains infected with two different *AR* silencing lentiviruses versus virus control with antibodies against phospho-Ser22 lamin A/C (pSer22; red) and DAPI for nuclear staining. Phospho-Ser22 lamin A/C fluorescence signal intensity assessed by Image J and expressed in arbitrary units (AU). n= 3 strains (>50 cells per condition), **p<0.01, non-parametric one-way ANOVA Kruskal–Wallis test, mean ± SE.

**Figure S5** (related to Fig. 5)

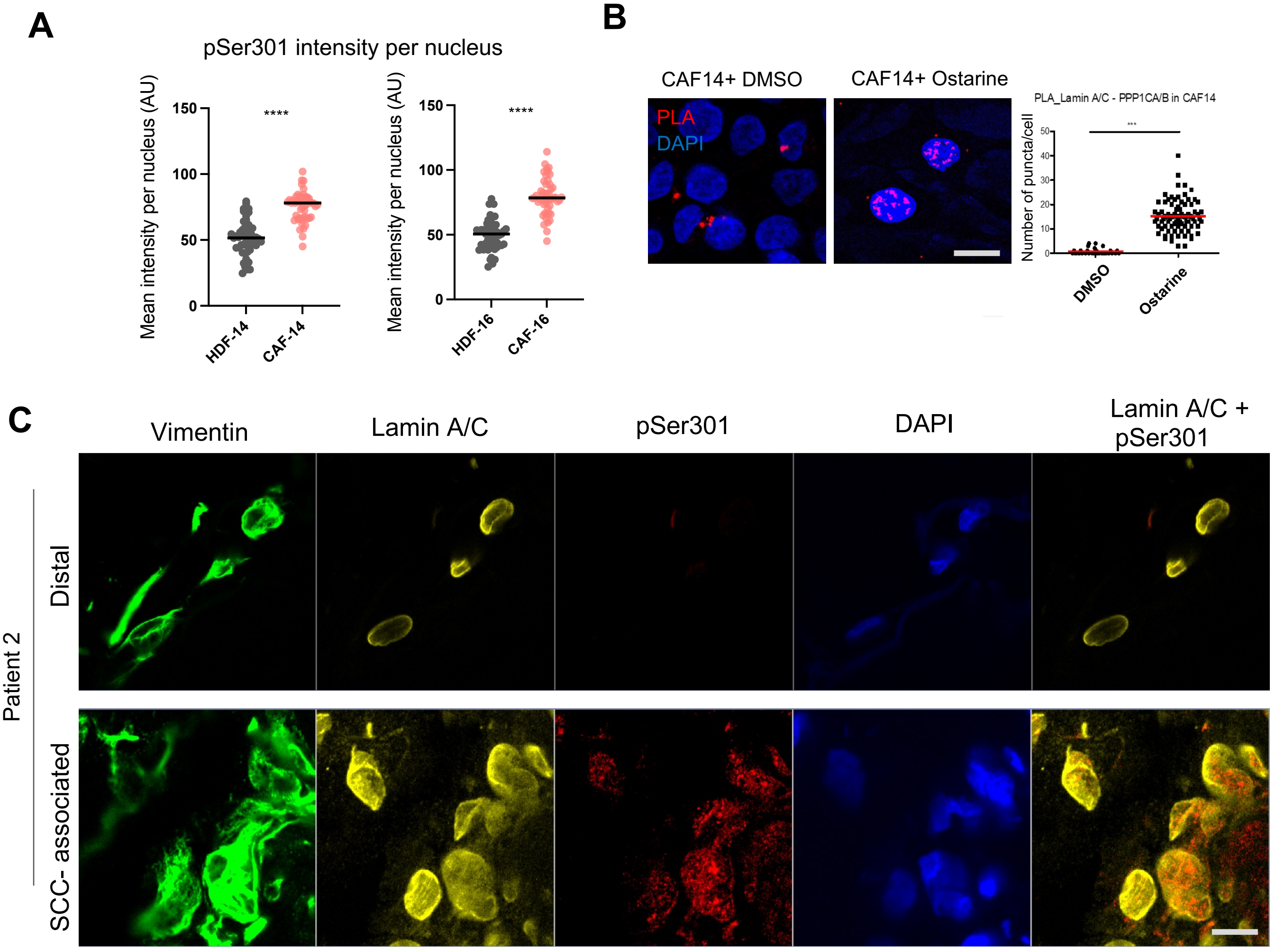


**Figure S5. Increased lamin A/C phosphorylation at ser 301 is a signature of CAFs.**

A) Quantification of pSer301 lamin A/C (pSer301) IF signal intensity in two pairs of CAFs and matched normal fibroblasts (HDFs) in addition to that shown in Fig. 5A. n(cells)>50 per condition, ****p<0.0001, unpaired student t-test. Scale bar: 10 μm.

B) Representative images and quantification of PLA signal for lamin A/C and PPP1CA/B association in CAFs treated with AR agonist ostarine (10 μM) or DMSO solvent control. n(cells)>50 per condition, ***p<0.001, unpaired student t-test.

C) Immunofluorescence analysis of a second skin SCC lesions and distal normal skin, beside that shown in Fig. 5D, with antibodies against pSer301-lamin A/C (red), total lamin A/C (yellow), anti-Vimentin (green; fibroblast marker), anti-Pan keratin (orange; epithelial marker).

### Figure S6 (related to Fig 6)

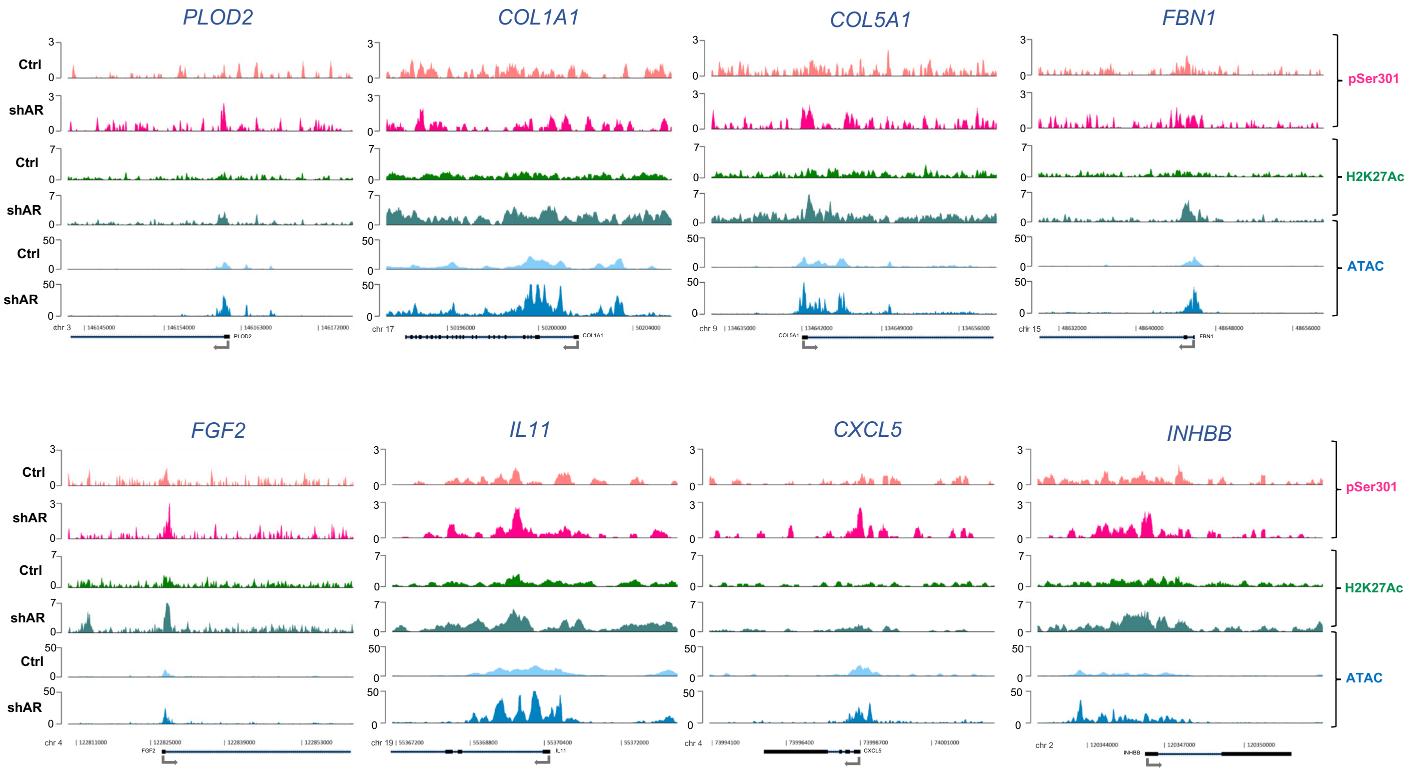


**Figure S6. Phospho-Ser301 lamin A/C binding to regulatory regions of CAF effector genes**

Illustration of pSer301-lamin A/C binding peaks on promoters/regulatory regions of selected CAF effector genes in control (Ctrl) and *AR* silenced (shAR) HDFs, beside those shown in Fig. 6E. ATAC-seq and H2K27Ac ChIP-seq profiles are used to identify open chromatin regions and promoter/enhancers, respectively. Position of the TSS and transcribed region is indicated.

### **Figure S7**. (related to Fig 7)

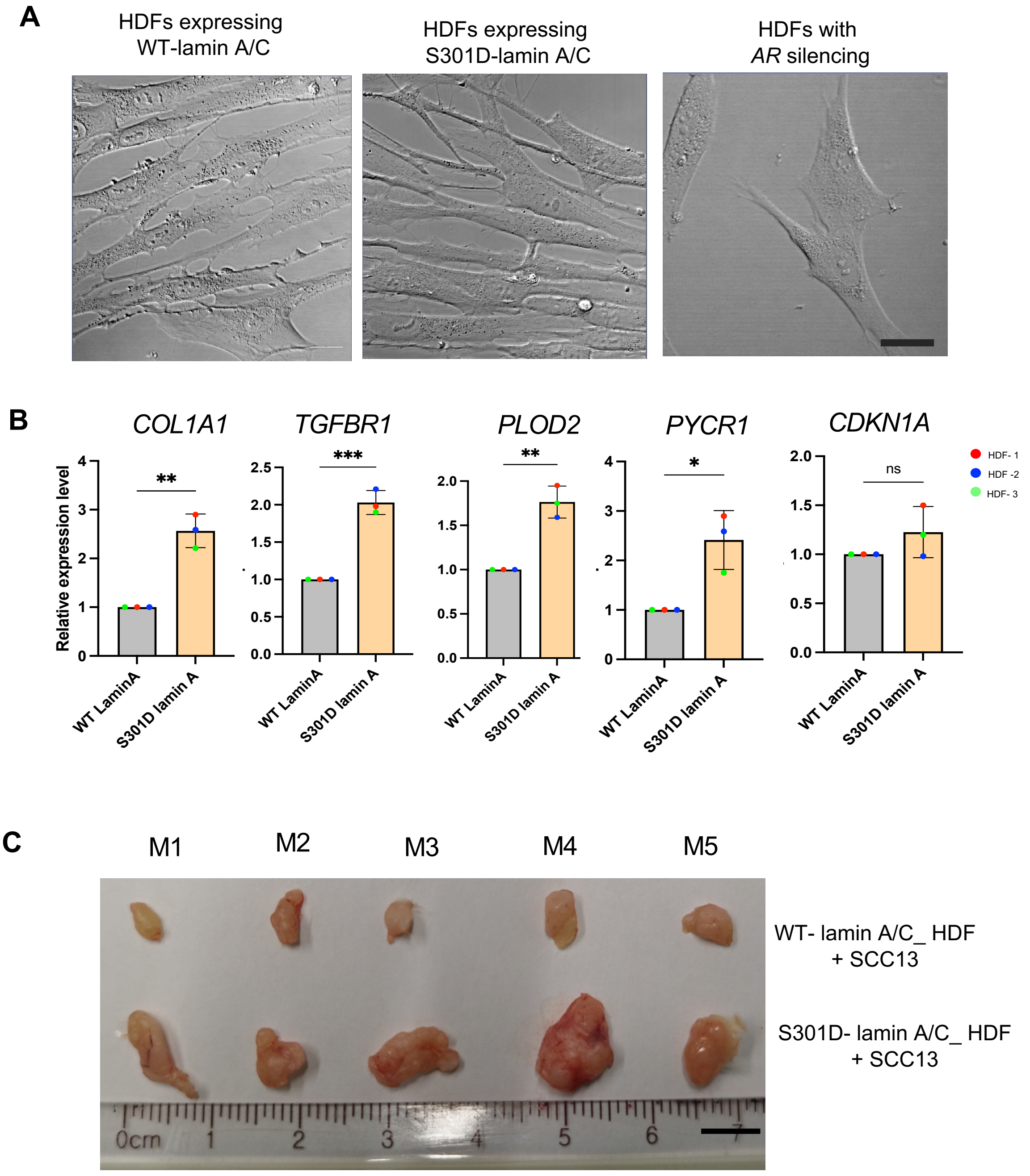


**Figure S7. Expression of a phosphomimetic Ser 301 lamin A/C mutant is sufficient to trigger CAF activation**.

A) Representative bright field images of HDFs transduced with lentiviruses expressing the S301D mutant or wild-type lamin A/C in parallel with cells with *AR* silencing. Scale bar: 20 µm.

B) RT-qPCR analysis of indicated genes in three different strains expressing S301D mutant or wildtype lamin A/C. Values are expressed relative to wildtype. n= 3 strains, *p<0.05,** p<0.005, ***p<0.001, unpaired t-test, mean ± SE.

C) Tumor formation assay. Images of excised xenograft tumor nodules formed by RFP expressing SCC13 cells co-injected intradermally with HDFs, expressing S301D mutant or wildtype lamin A/C, in contralateral mouse back skin. Quantification of the tumor volume is shown in Fig.7E. Scale bar is 5 mm.

#### Table S1. List of reagents and resources used in this study

| REAGENT or RESOURCE | | |
|---|---|---|
| **Antibodies** | SOURCE | Catalog number |
| Androgen Receptor (D6F11) XP Rabbit mAb antibody | Cell Signaling Technology | Cat# 5153S |
| Rabbit Anti-Androgen Receptor, PG-21 Polyclonal antibody, Unconjugated | Millipore | Cat# 06-680 |
| β-Actin (8H10D10) Mouse mAb antibody | Cell Signaling Technology | Cat# 3700 |
| Mouse Anti-gamma-Tubulin Monoclonal Antibody, Unconjugated, Clone GTU-88 | Sigma-Aldrich | Cat# T6557 |
| Rabbit Anti-Androgen Receptor Polyclonal Antibody, Unconjugated | Abcam | Cat# ab74272 |
| Lamin A/C (4C11) Mouse mAb | Cell Signaling Technology | Cat# 4777 |
| Lamin A antibody + Lamin C antibody [EPR4100] | Abcam | Cat# ab108595 |
| Phospho-Lamin A/C (Ser301) Antibody | Affinity Biosciences | Cat# AF7177 |
| Histone H3 (acetyl K27) antibody - ChIP Grade | Abcam | Cat# ab4729 |
| Rabbit Anti-Histone H3, trimethyl (Lys9) ChIP Grade Polyclonal Antibody, Unconjugated | Abcam | Cat# ab8898 |
| Anti-Lamin B1 antibody [EPR8985(B)] | Abcam | Cat# ab133741 |
| GAPDH Antibody (FL-335) | Santa Cruz Biotechnology | Cat# sc-25778 |
| Rat monoclonal anti-Vimentin (Clone # 280618) | R&D Systems | Cat# MAB2105 |
| Anti-Cytokeratin Pan Antibody, clone Lu5 | Acris Antibodies | Cat# BM4068 |
| Lamin A/C (636) antibody | Santa Cruz Biotechnology | Cat# sc-7292, |
| Anti-Rabbit IgG (H+L), HRP Conjugate antibody | Promega | Cat# W401B |
| Anti-Mouse IgG (H+L) Antibody, HRP Conjugated | Promega | Cat# W4021 |
| Donkey Anti-Mouse IgG (H+L) Antibody, Alexa Fluor 488 Conjugated | ThermoFisher | Cat# A-21202 |
| Donkey anti-Rabbit IgG (H+L) Highly Cross-Adsorbed Secondary Antibody, Alexa Fluor 568 | ThermoFisher | Cat# A-10042 |
| Donkey anti-Goat IgG (H+L) Cross-Adsorbed Secondary Antibody, Alexa Fluor™ 647 | ThermoFisher | Cat# A-21447 |
| Chicken anti-Rat IgG (H+L) Cross-Adsorbed Secondary Antibody, Alexa Fluor™ 647 | ThermoFisher | Cat# A-21472 |
| Donkey anti-Rabbit IgG (H+L) Highly Cross-Adsorbed Secondary Antibody, Alexa Fluor™ Plus 488 | ThermoFisher | Cat# A-21206 |
| Donkey anti-Mouse IgG (H+L) Highly Cross-Adsorbed Secondary Antibody, Alexa Fluor™ 568 | ThermoFisher | Cat# A10037 |
| Chicken anti-Rabbit IgG (H+L) Cross-Adsorbed Secondary Antibody, Alexa Fluor™ 647 | ThermoFisher | Cat# A-21443 |
| Donkey anti-Goat IgG (H+L) Cross-Adsorbed Secondary Antibody, Alexa Fluor™ 488 | ThermoFisher | Cat# A-11055 |

| **Biological Samples** | SOURCE | IDENTIFIER |
|---|---|---|
| HDFs: Foreskin and abdominoplasty skin samples from Lausanne University Hospital | CHUV, Lausanne, Switzerland | N/A |
| CAFs: surgically excised skin SCCs from Massachusetts General Hospital | Department of Dermatology, Massachusetts General Hospital (Boston, MA) | N/A |
| Frozen human tissue blocks from matched normal and SCC skin | Department of Dermatology, Massachusetts General Hospital (Boston, MA) | N/A |

| **Special Chemicals** | SOURCE | Catalog number |
|---|---|---|
| 5α-Androstan-17β-ol-3-one / Dihydrotestosterone | Sigma-Aldrich | Cat# 10300 |
| AZD3514 | AdooQBiosciences | Cat# A12396 |
| Ostarine | Selleckchem | Cat# S1174 |
| ARCC4 | Tocris | Cat# 7254 |
| Tautomycin | Santa Cruz | Cat# sc-507214 |
| Matrigel (Growth Factor Reduced) | Corning | Cat# 354230 |

| **Critical Commercial Assays** | SOURCE | Catalog number |
|---|---|---|
| SuperSignal™ West Pico PLUS Chemiluminescent Substrate | Thermo Fisher Scientific | Cat# 34578 |
| Agilent RNA 6000 Nano Kit | Agilent | Cat# 5067-1511 |
| Direct-zol RNA Miniprep Kit | Zymo Research | Cat# R2050 |

| **Experimental Models: Organisms/Strains** | | |
|---|---|---|
| Mouse: NOD.Cg-Prkdcscid Il2rγtm1Wjl/SzJ | The Jackson Laboratory | JAX:005557 |

| **Experimental Models: primary cells and cell lines** | SOURCE | IDENTIFIER |
|---|---|---|
| HDF-1 | CHUV, Lausanne, Switzerland | AK-1 |
| HDF-2 | CHUV, Lausanne, Switzerland | PB-2 |
| HDF-3 | CHUV, Lausanne, Switzerland | AK-2 |
| HDF-11 | Clocchiatti, A. *et al.* J Clin Invest 2018 | N/A |
| CAF-11 | Clocchiatti, A. *et al.* J Clin Invest 2018 | N/A |
| HDF-14 | Clocchiatti, A. *et al.* J Clin Invest 2018 | N/A |
| CAF-14 | Clocchiatti, A. *et al.* J Clin Invest 2018 | N/A |
| HDF-14 | Clocchiatti, A. *et al.* J Clin Invest 2018 | N/A |
| CAF-14 | Clocchiatti, A. *et al.* J Clin Invest 2018 | N/A |
| SCC13 | Restivo, G. et al. EMBO J, 2011 | CVCL_4029 |
| **sgRNA details** | SOURCE | IDENTIFIER/ Sequences |
| ctrl gRNA | ABM Bio | Cat# K011 |
| sgRNA AR KO-1 | ABM Bio | 5'-ATTCTGGAAAGCTCCTCGGT-3' |
| sgRNA ARKO-2 | ABM Bio | 5'-CGCGAAGTGATCCAGAACCC -3' |

| **Recombinant DNA** | SOURCE | IDENTIFIER |
|---|---|---|
| pLKO.1shAR 1 (human) | Sigma-Aldrich | TRCN0000003718 |
| pLKO.1shAR 2 (human) | Sigma-Aldrich | TRCN0000003715 |
| pLenti-lacZ | Clocchiatti, A. et al. J Clin Invest 2018 | N/A |
| pLenti-AR-V5 | Azeem W et al, 2017 | N/A |
| pCDH_blast_MCS_Nard_GFP_LAMIN | Addgene | Cat# 167340 |
| pCDH_blast_MCS_Nard_GFP_LAMIN_S301D | This study | N/A |
| pTRIP-SFFV-EGFP-NLS | Addgene | Cat# 86677 |
| **Software and Algorithms** | SOURCE | IDENTIFIER |
| GSAA-seqSP, version GSAA_2.0 | Xiong et al., 2014 | http://gsaa.unc.edu |
| R package 'survival' v2.43-3 | Terry M et al, 2000 | https://CRAN.R-project.org/package=survival |

| **RT-PCR primers (human and mouse)** | |
|---|---|
| **Gene ID** | **Primer sequence (5'-3')** |
| *RPLP0* | Forward: GCAATGTTGCCAGTGTCTGT |
| | Reverse: GCCTTGACCTTTTCAGCAAG |
| *AR* | Forward: TTGTGTCAAAAGCGAAATGG |
| | Reverse: AGTCAATGGGCAAAACATGG |
| *CDKN1A* | Forward: CCCAAGCTCTACCTTCCCAC |
| | Reverse: ACAGGTCCACATGGTCTTCC |
| *PLOD2* | Forward: GAACAGCCCTGTCCAGATGT |
| | Reverse: CATGATGTTTCCCCCAGAC |
| *COL1A1* | Forward: GATTCCCTGGACCTAAAGGTGC |
| | Reverse:AGCCTCTCCATCTTTGCCAGCA |
| *TGFBR1* | Forward: GAGCATGGATCCCTTTTTGA |
| | Reverse: ATGTGAAGATGGGCAAGACC |
| *PYCR1* | Forward: CCCCGCCTACGCATTCACA |
| | Reverse: GCGCGTTGGAAGTCCCATCT |
